# MeCP2 regulates cell type-specific functions of depressive-like symptoms in the nucleus accumbens

**DOI:** 10.1101/2025.07.24.666523

**Authors:** Jinhee Bae, Sung Hoon Kim, Sun A Jung, Nazarii Frankiv, Eun Mi Hwang, Young-Min Kim, Sangjoon Lee, Heh-In Im

**Author notes:** Corresponding author: H.-I. Im.

## Abstract

MeCP2 (methyl CpG binding protein 2) is a transcriptional regulator that modulates gene expression in response to environmental stimuli. Although recent studies have implicated MeCP2 in stress responses and depression, its precise role is not completely understood. In this study, we identify a cell type-specific function of MeCP2 in the regulation of depression-like symptoms within the nucleus accumbens (NAc), a key brain region for emotional and stress processing. We observed differential MeCP2 expression in distinct cell populations of the NAc following chronic restraint stress (CRS) and investigated the behavioral and electrophysiological consequences of cell type-specific MeCP2 manipulation. We also explored the molecular mechanisms by which MeCP2 alleviates depression-like symptoms in the NAc and associated neural circuit regions through cell type-specific profiling of the spatial transcriptome. Our findings demonstrate that MeCP2 contributes to synaptic and circuit-level regulation in a cell type-specific manner within the NAc and ultimately mitigates CRS-induced depression-like behaviors.

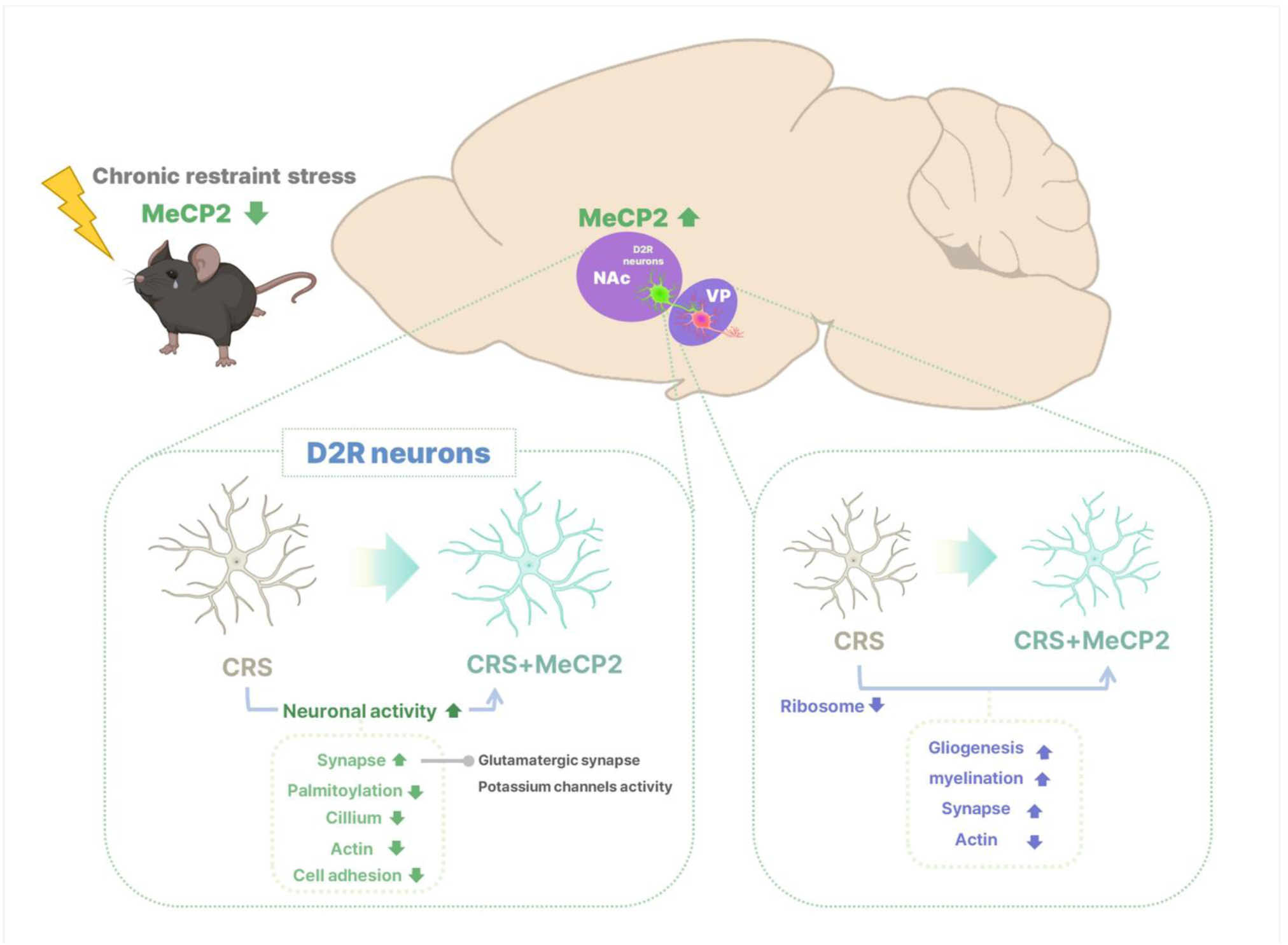
Graphical abstract. Schematic Summary of the functional role of accumbal MeCP2

## Introduction

MeCP2 (methyl CpG binding protein 2) is a transcriptional regulator that modulates gene expression in response to environmental stimuli(Chahrour, Jung *et al*., 2008; Guy, Cheval *et al*., 2011), including stress and drug exposure (Im, Hollander *et al*., 2010; Su, Hong *et al*., 2015; Lewis, Bastle *et al*., 2016; Sánchez-Lafuente, Kalynchuk *et al*., 2022). While MeCP2 is best known for its role in neurodevelopmental disorders, accumulating evidence suggests that it also plays a critical role in stress adaptation and the pathophysiology of depression(Su, Hong et al., 2015; Lewis, Bastle et al., 2016; Huang, He *et al*., 2022). For instance, phosphorylation of MeCP2 at serine 421 in the hippocampus has been shown to be essential for the sustained antidepressant effects of ketamine in rodents^3^ and decreased MeCP2 protein levels have been reported in the blood of patients with major depressive disorder(Su, Hong et al., 2015). In addition, differential expression of MeCP2 has been observed between stress-resilient and susceptible mice following chronic social defeat stress, implicating its involvement in stress vulnerability(Huang, He et al., 2022). Despite these findings, the precise role of MeCP2 in regulating depressive symptoms remains largely unexplored. In particular, the neurophysiological mechanisms through which MeCP2 modulates affective behavior under stress conditions are not well understood. Addressing this knowledge gap is essential to advance our understanding of the biological mechanisms underlying stress-related mood disorders.

The nucleus accumbens (NAc) is one of the core regions of the reward circuitry and plays a critical role in stress responses and mood regulation (Russo and Nestler, 2013; Francis and Lobo, 2017; Jiang, Zou *et al*., 2023). Although recent studies have suggested that MeCP2 in the NAc may contribute to stress-related processes(Lewis, Bastle et al., 2016), the biological mechanisms through which MeCP2 regulates stress or depressive symptoms in this region remain poorly understood. In particular, the two cell types that dominantly compose the NAc, dopamine 1 receptor (D1R) and dopamine 2 receptor (D2R) neurons, are known to contribute differently to stress, depression, reward, and motivation regulation(Nestler and Carlezon Jr, 2006; Bewernick, Hurlemann *et al*., 2010). Given the functional heterogeneity of these cell types, a cell type–specific approach is essential to elucidate the gene-specific mechanisms underlying stress and affective disorders in the NAc.

In this study, we found that chronic restraint stress (CRS) downregulated MeCP2 in the NAc specifically in D2R neurons whereas MeCP2 overexpression in these neurons alleviated depressive-like symptoms. To elucidate the underlying neurobiological mechanisms, we compared the neural activity and transcriptomic profiles across control, diseased (CRS-exposed model), and rescued (a model combining CRS-exposed and MeCP2-rescued) groups. Notably, MeCP2 manipulation in the NAc not only reversed CRS-induced behavioral symptoms but also rescued CRS-altered gene expression patterns at the circuit level. These findings suggest that MeCP2 regulates depressive-like behaviors in the NAc via cell type–specific molecular and physiological pathways.

## Material and Methods

Comprehensive descriptions of all experimental protocols are provided in the Supplementary Infromation.

## Results

### Chronic restraint stress (CRS) induces depressive-like behavior and reduces MeCP2 only in D2R neurons in the NAc

To confirm the association of depression and MeCP2 in the NAc, we first exposed C57BL/6J male mice to chronic restraint stress (CRS) (Fig. 1a). Except for immediately before CRS exposure (day 1), CRS group mice showed significant decreases in body weight compared to the control group on all other days (Fig. 1b). CRS did not change the distance traveled by mice in an open field test (Fig. 1c, d), indicating that it did not impair the general motor function of mice. CRS also did not decrease the time spent in the center zone of an open field box, a measure of anxiety level (Fig. 1e), but decreased the time spent in the open arm of an elevated plus maze (Figs. 1f, g). In addition, it increased immobility in a forced swim test (Figs. 1h, i), indicating that CRS induced anxiety-like and depressive-like expressions.

**Fig. 1.**
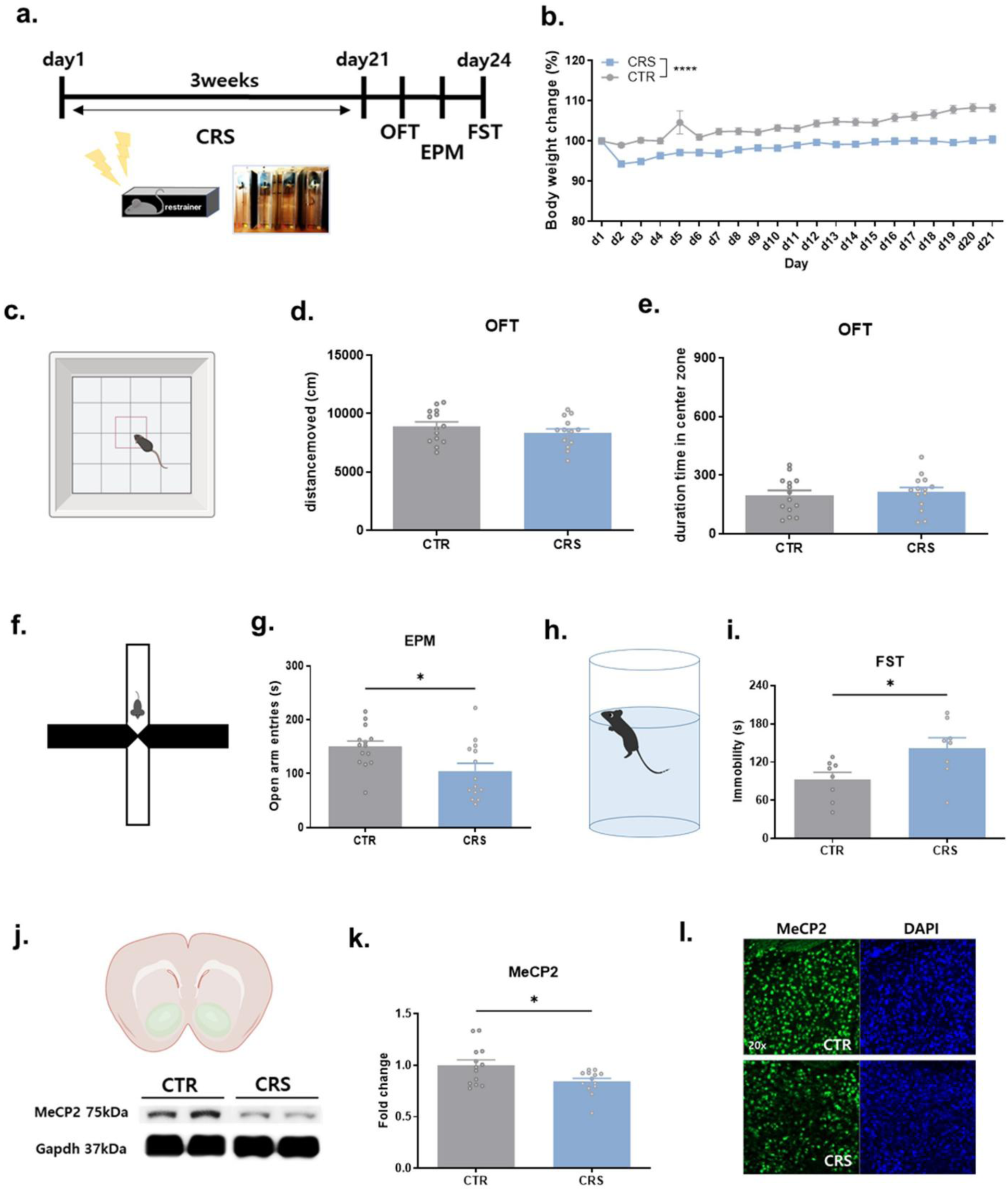
Effect of chronic restraint stress (CRS) in NAc **a** Timeline of the experiment. OFT: open field test, EPM: elevated plus maze FST: forced swim test **b** Changes in body weight over 3 weeks of exposure to CRS. CTR: control. Two-way ANOVA mixed-model, F (20, 600) = 4.330, p < 0.0001 (n=16/group) **c-e** Open field test. CRS did not affect locomotion (t-test, two-tailed, t = 1.111, p = 0.2767, df = 26, n=14 mice/group) or time spent in the center zone (t-test, two-tailed, t = 0.4432, p = 0.6613, df = 26, n=14 mice/group). **f-i** CRS increased anxiety and depressive symptoms**. f, g** Elevated plus maze test (t-test, two-tailed, t = 2.561, p = 0.0166, df = 26, n =14 mice/group) **h, i** Forced swim test (t-test, two-tailed, t = 2.498, p = 0.0256, df = 14, n = 8 mice/group) **j-l** Changes in MeCP2 protein levels in the NAc by CRS **j, k** Western blot detection of MeCP2 in the control and CRS groups **j** Representative immunoblots in each group **k** Quantification of Western blot bands (t-test, two-tailed, t = 2.537, p = 0.0181, df = 24, n = 13 mice/group) **l** Representative immunofluorescence images showing the decrease of MeCP2 in the NAc (20X magnification

We investigated whether CRS exposure changes the expression level of MeCP2 protein in the NAc. We observed a significant decrease in the expression level of MeCP2 protein in the NAc of mice three days after the last CRS exposure compared to the control group (Figs. 1j-l). The expression of MeCP2 was not changed in the dorsal striatum, medial prefrontal cortex, or hippocampus, which are involved in motivational and emotional regulation (Supplementary Fig. 1). The same decrease in MeCP2 was observed 2 h after the last CRS (Supplementary Fig. 2).

Recent studies have reported that D1R and D2R neurons, two types of cells that make up the majority of NAc neurons, regulate depressive symptoms differently and that related genes are changed differently in each cell type in depression models. Therefore, we examined whether the expression level of MeCP2 in the NAc by CRS shows different patterns depending on cell type. We first distinguished the two cells using antibodies reactive to D1 and D2 receptors (Supplementary Figs. 3a, b), and confirmed the expression changes in each cell type in the NAc of CRS-exposed mice. Interestingly, CRS exposure significantly reduced MeCP2 in D2R neurons but not in D1R neurons (F2a-c, Supplementary Figs. 4a, b). This difference in expression was the same 2 h after the end of CRS (Supplementary Figs. 5a-e).

### MeCP2 regulates depressive phenotypes in a cell type-specific manner in the NAc

To investigate the association between the reduction of MeCP2 in D2R neurons in the NAc and the induction of depressive symptoms, we constructed cell type-specific AAV vectors expressing shMeCP2 only in D2 neurons (Fig. 2d-g). AAV-G-CREon-scrambled or AAV-G-CREon-shMeCP2 was injected bilaterally into the NAc of D2Cre mice. The reduction of MeCP2 protein and mRNA levels was confirmed by immunohistochemistry and qPCR from the cells isolated by fluorescence-activated cell sorting (FACS) (Figs. 2e-g, Supplementary Fig. 6).

**Fig. 2.**
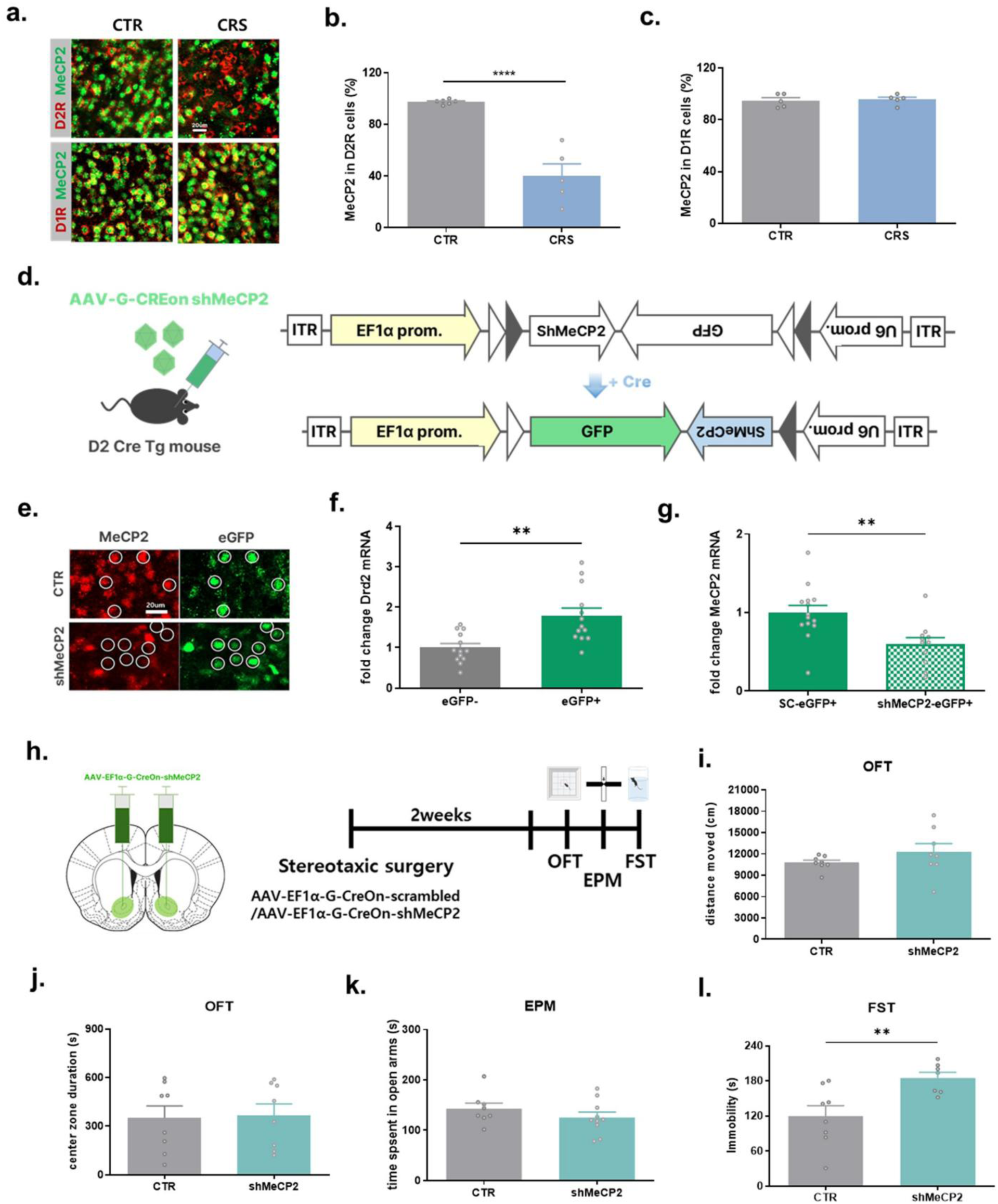
MeCP2 regulates CRS in a cell-type-specific manner in the NAc. **a-c** CRS reduces MeCP2 in D2R neurons. **a** Representative immunofluorescence of MeCP2 and D1R receptor/D2R receptors in the NAc at 3 days after CRS termination (scale bar: 20um) **b, c** Quantification graphs of the number of MeCP2 positive cells among D1R/D2R receptor positive cells **b** MeCP2 expression in D2R neuron ( t-test, two-tailed, t = 6.741, p <0.0001, df = 9, n = 6, 5 mice/group) **c** MeCP2 expression in D1R neuron ( t-test, two-tailed, t = 0.3253, p = 0.7533, df = 8, n = 5 mice/group) **d** Diagrams for G-CREon viral vector containing MeCP2 shRNA treated by CRE recombinase **e-f** Functional validation of AAV-G-CREon shMeCP2. **e** Representative images showing the reduction of MeCP2 protein. CTR (control): AAV inserted with scrambled-shRNA. **f, g** Characterization of cells isolated by FACs sorting. **f** Higher expression level of Drd2 mRNA in eGFP-positive cells than in eGFP-negative cells (eGFP- vs. eGFP+: paired t-test, two-tailed, t = 3.781, p = 0.0026, df = 12, n = 13 mice/group). **g** Reduced expression level of MeCP2 mRNA in cells expressing MeCP2 shRNA compared to scrambled shRNA (SC) (SC-eGFP+ vs. shMeCP2-eGFP+: t-test, two-tailed, t = 3.136, p = 0.0046, df = 23, n = 14, 11 mice/group). **h** Experimental schedule to confirm the function of shMeCP2 in depression. MeCP2 was reduced in a cell type-specific manner using D2R Cre mice. **i-l** In D2R neurons, shMeCP2 increased depressive symptoms (l) but did not affect locomotion (i) and anxiety levels (j,k). **i** Total distance moved in OFT (t-test, two-tailed, t = 1.219, p = 0.2429, df = 14, n = 8 mice/group) **j** Time spent in center zone in OFT (t-test, two-tailed, t = 0.1585, p = 0.8763, df = 14, n = 8 mice/group) **k** Time spent in open arms in EPM (t- test, two-tailed, t = 1.131, p = 0.2747, df = 16, n = 8, 10 mice/group)**ㅣ** Immobility time in FST (t-test, two-tailed, t = 3.073, p = 0.0089, df = 13, n = 8, 7 mice/group) All data are shown as mean ± SEM. **p <0.01, **** p <0.0001

We confirmed that MeCP2 knockdown in D2R neurons without exposure to CRS could induce depressive-like symptoms. shMeCP2 mice showed significantly increased immobility in a forced swim test compared to the control group. This suggests that MeCP2 knockdown in NAc D2R neurons could be a major cause of depressive symptoms (Fig. 2l). In contrast, MeCP2 knockdown in D2R neurons did not affect locomotion or anxiety levels (Figs. 2i-k).

Next, to determine whether restoring MeCP2 in D2R neurons where it had been reduced by CRS has a therapeutic effect on depression, we constructed a vector expressing MeCP2 in a cell type-specific manner (Fig. 3a). The increased expression of MeCP2 protein by viral manipulation was confirmed using immunohistochemistry (Fig. 3b, Supplmentary Fig. 7). AAV-EF1alpha-DIO-eGFP or AAV-EF1alpha-DIO-mMeCP2-eGFP was bilaterally injected into the NAc of D2Cre mice. Two weeks after virus injection, the mice in the treatment group were exposed to CRS for three weeks (Fig. 3c). Interestingly, the CRS exposure group with increased MeCP2 expression underwent a significant increase in body weight compared to the CRS group, although their weight did not fully recover to the levels observed in the naïve control group (Fig. 3d). In addition, depressive symptoms were improved to the naive-control level in elevated plus maze and forced swim tests (Fig. 3g, h).

**Fig. 3.**
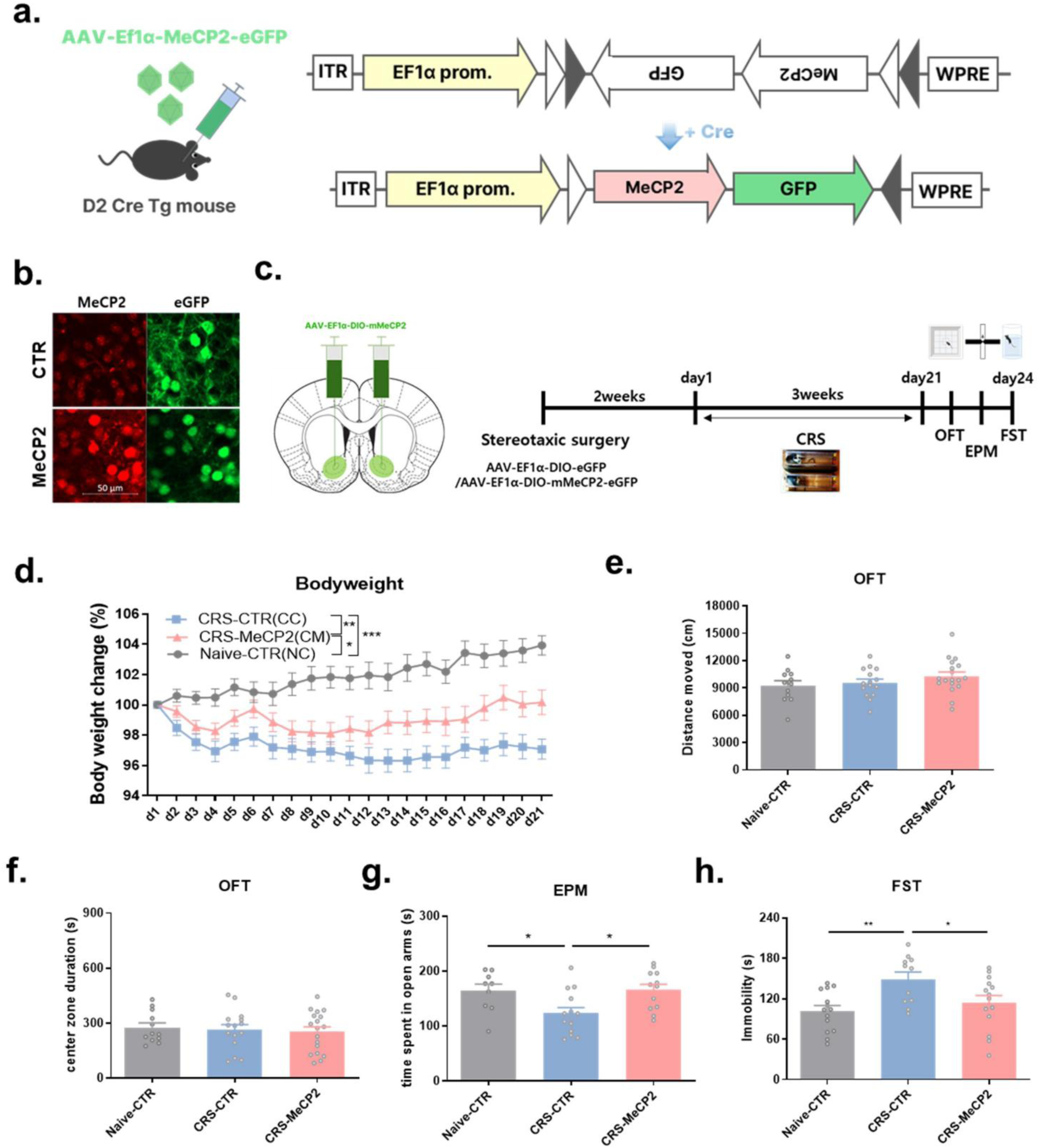
Alleviation of depressive symptoms by genetic MeCP2 upregulation in D2R neurons in the NAc **a** Diagram of viral vectors that increase cell type-specific mouse MeCP2 mRNA by CRE recombinase **b** Increased MeCP2 protein in cells expressing AAV-MeCP2. CTR: control-virus. Scale bar: 50 μm **c** Experimental procedures to confirm the therapeutic efficacy of genetic MeCP2 augmentation in the CRS model. **d** Graph of body weight change (%) during the CRS exposure period. Two-way Mixed ANOVA, post-hoc Holm-Šídák’s test, n = 13(NC), 19(CC), 19(CM) mice/group, NC vs. CC: p <0.001, NC vs. CM: p = 0.003, CC vs CM: p = 0.019 **e** Distance moved in OFT (One-way ANOVA, F (2, 42) = 1.357, p = 0.2685, post-hoc Holm-Šídák’s test, NC vs.CC: p = 0.6702, NC vs. CM: p = 0.3404, CC vs CM: p = 0.4334, n=12, 15, 18 mice/group) **f** Time spent in center zone in OFT, One-way ANOVA, F (2, 42) = 0.1852, p = 0.8316, post-hoc Holm-Šídák’s test, NC vs.CC: p = 0.9425, NC vs. CM: p = 0.9074, CC vs CM: p = 0.9425) **g** Time spent in open arms in EPM (One-way ANOVA, F (2, 32) = 5.241, p = 0.0107, post-hoc Holm-Šídák’s test, NC vs.CC: p = 0.0320, NC vs. CM: p = 0.9036, CC vs. CM: p = 0.0201, n = 9, 14, 12 mice/group) **h** immobility time in FST (One-way ANOVA, F (2, 36) = 5.570, p = 0.0078, post-hoc Holm-Šídák’s test, NC vs.CC: p = 0.0072, NC vs. CM: p = 0.3587, CC vs CM: p = 0.0434, n = 14, 11, 14 mice/group). All data are shown as mean ± SEM. * p <0.05, **p <0.01, p <0.001 ***

Upregulation of MeCP2 in D2R neurons did not affect the locomotion and center zone time of mice in the CRS exposure group (Figs. 3e, f). In addition, upregulation of MeCP2 expression in D2R neurons of normal mice did not induce cognitive or motor dysfunction, indicating that the increase of MeCP2 in D2R neurons in NAc did not cause negative side effects on general functions (Supplementary Fig. 8). Additionally, we investigated whether MeCP2 overexpression in D1R neurons could modulate depressive symptoms induced by CRS exposure. However, upregulation of MeCP2 in D1R neurons in NAc did not alleviate CRS-induced depressive symptoms (Supplementary Fig. 9). This suggests that upregulation of MeCP2 only functions to alleviate depressive symptoms in D2R neurons. Taken together, our results demonstrate that repeated and persistent stress induces a decrease in D2R neuron-specific MeCP2 in the NAc and that cell type-specific regulation of MeCP2 in the NAc may contribute to alleviating depression.

### MeCP2 overexpression in D2R neurons rescues the altered neural activity after CRS

Our findings, including in particular a significant decrease in MeCP2 in D2R neurons in the NAc after CRS exposure and changes in depressive symptoms by manipulating expression of MeCP2, demonstrate the functional importance of MeCP2 in depression. Therefore, we investigated the neurophysiological mechanism by which MeCP2 in D2R neurons exerts therapeutic efficacy. To elucidate the neurological mechanism underlying MeCP2 regulation of depression in D2R neurons of the NAc, we first investigated whether genetic manipulation of MeCP2 induces changes in neuronal activity in depression. Changes of neuronal activity in the brain are known to be a representative symptom of depression(Marsden, 2013; Fries, Saldana *et al*., 2023). Based on recent reports on the function of MeCP2 in regulating neuronal activity(Huang, Zhang *et al*., 2021; Kim, Autry *et al*., 2021), we hypothesized that the level of neuronal activity would be altered in the NAc of the CRS model in which MeCP2 was reduced. In addition, we investigated whether MeCP2 upregulation rescues the changes in neural activity induced by CRS. To confirm this, we measured electrophysiological responses in ex vivo nucleus accumbens slices using a multielectrode array (MEA) recording system and optogenetics. We co-expressed MeCP2 and channelrhodopsin in D2R neurons of the NAc by injecting AAV viruses expressing cell types in a cell-type specific manner into D2R Cre mice (Figs. 4a-c, Supplementary Fig. 10a). In response to optical stimulation (473 nm, 20 pulses of 25 ms for 1 s at 20 Hz, 50% duty cycle; 10 mW at the plate bottom), the CRS group showed significantly decreased neural activity in the NAc compared with the control group. This suggests that CRS exposure changed the neural activity level of D2R neurons (Figs. 4b, d–f, Supplementary Fig. 10b). In eight repeated sessions, the response pattern of neural activity to optical stimulation was observed to be the same, without any increase or decrease (Fig. 4e). Overexpression of MeCP2 significantly restored the altered neural activity in the NAc after CRS exposure.

**Fig. 4.**
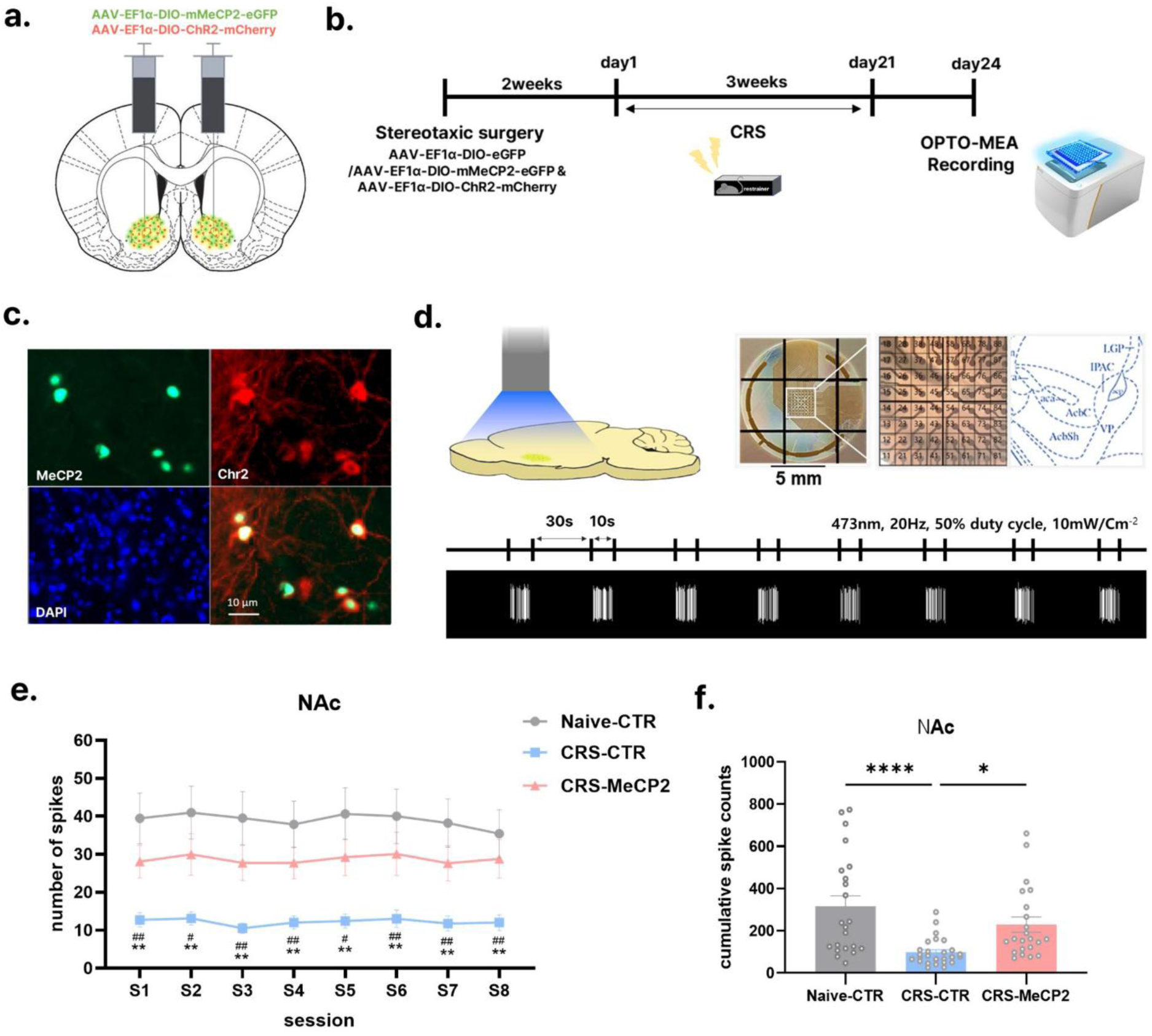
Modulation of neural activity by genetic MeCP2 augmentation **a** AAV injection inducing cell type-specific coexpression of MeCP2 and Chr2 in the NAc. **b** Experimental schedule to confirm the modulation of neural activity by genetic MeCP2 augmentation in a depression model. **c** Representative immunofluorescence images showing the expression of MeCP2 and Chr2. Scale bar: 10 μm. **d** Electrophysiological measurements using optogenetics and multielectrode array (MEA) recording. A sagittal slice of the NAc is placed on a 64-channels MEA recording plate (upper right panel). Blue light was presented for 10 s with a 30-s interval. A total of eight stimuli were given. **e** Total number of spikes for each 10-s session during the optogenetic stimulation. S1-S8(session1-session8), Naive-CTR(NC), CRS-CTR(CC), CRS-MeCP2(CM), Two-way Mixed ANOVA, F(2, 68) = 10.47, p = 0.0001, n = 22, 27, 22 mice/ group, post-hoc Holm-Šídák’s test, S1: NC vs. CC p = 0.0022, NC vs. CM p = 0.1605, CC vs. CM p= 0.0055, S2: NC vs. CC p = 0.0022, NC vs. CM p = 0.2212, CC vs. CM p = 0.0143, S3: NC vs. CC p = 0.0015, NC vs. CM p = 0.1665, CC vs. CM p = 0.0028, S4: NC vs. CC p = 0.0013, NC vs. CM p = 0.1790, CC vs. CM p = 0.0035, S5: NC vs. CC p = 0.0016, NC vs. CM p = 0.1795, CC vs. CM p = 0.0059, S6: NC vs. CC p = 0.0042, NC vs. CM p = 0.2853, CC vs. CM p = 0.0189, S7: NC vs. CC p= 0.0016, NC vs. CM p = 0.1851, CC vs. CM p = 0.0073, S8: NC vs. CC p = 0.0049, NC vs. CM p = 0.4170, CC vs. CM p = 0.0094 #p <0.05, ##p <0.01 (CC vs. CM), *p <0.05, **p <0.01 (NC vs. CC) **f** Total cumulative spike counts across all sessions. One-way ANOVA, F(2, 68) = 10.56, p = 0.0001, post-hoc Holm-Šídák’s test, NC vs. CC p <0.0001, NC vs. CM 0.0937, CC vs. CM p = 0.0156, *p <0.05, ****p <0.0001. All data are shown as mean ± SEM.

### D1R and D2R neurons in the NAc function differently

Next, we identified the neurobiological molecular pathways by which MeCP2 in D2R neurons modulates depression. We explored the pathways by which MeCP2 modulates neural activity to alleviate depressive symptoms and investigated the transcriptomic networks of downstream genes regulated by MeCP2. Using GeoMx Digital Spatial Profiler (DSP) technology, we isolated D2R neurons from the NAc and performed a differential gene expression (DEG) analysis on the effects of CRS and MeCP2 regulation (Fig. 4a-c). In the DEG analysis, we focused on deriving genes among those altered by CRS that showed CRS effects rescued by MeCP2 overexpression and to elucidate their molecular and biological functions. Furthermore, we also investigated whether MeCP2 overexpression in the NAc resulted in molecular rescue at the circuit level (Fig. 4c). Building on the observation that overexpression of MeCP2 in the NAc alleviated depressive-like behaviors, we postulated that similar molecular recovery might be observed across anatomically and functionally connected brain regions. We selected the ventral pellidum, which is the region where most of the D2R neurons of the NAc project, as the target region because it is most directly affected by and primarily responds to changes in the NAc D2R neurons.

To selectively manipulate gene expression, D2R-expressing neurons in the NAc were transduced with AAV vectors encoding MeCP2 and eGFP or eGFP alone (Figs. 3a-c). Mice were then subjected to chronic restraint stress (CRS) for three weeks, resulting in three groups: control (Naive-control, NC), diseased (CRS-exposed model, CRS-control,CC), and rescued (a model combing CRS-exposed and MeCP2-rescuedCRS-MeCP2, CM) (Fig. 4a). Three days after the end of CRS, the mice brains were extracted and a spatial transcriptome analysis was performed using GeoMX DSP. For a cell type-specific analysis, fluorescence in situ hybridization for D1 and D2 dopamine receptors (Drd1, Drd2) and eGPF mRNAs was used to distinguish cell types in sagittal sections of the mouse brains (Figs. 5b-d, Supplementary Fig. 11). We confirmed that the two cell types were well distinguished through each marker (Supplementary Figs. 11b,c). In addition, we confirmed that the markers of eGFP and D2R neurons were coexpressed. This verifies that AAV functioned well in a cell type-specific manner (Fig. 5e, Supplementary Fig. 11d).

**Fig. 5.**
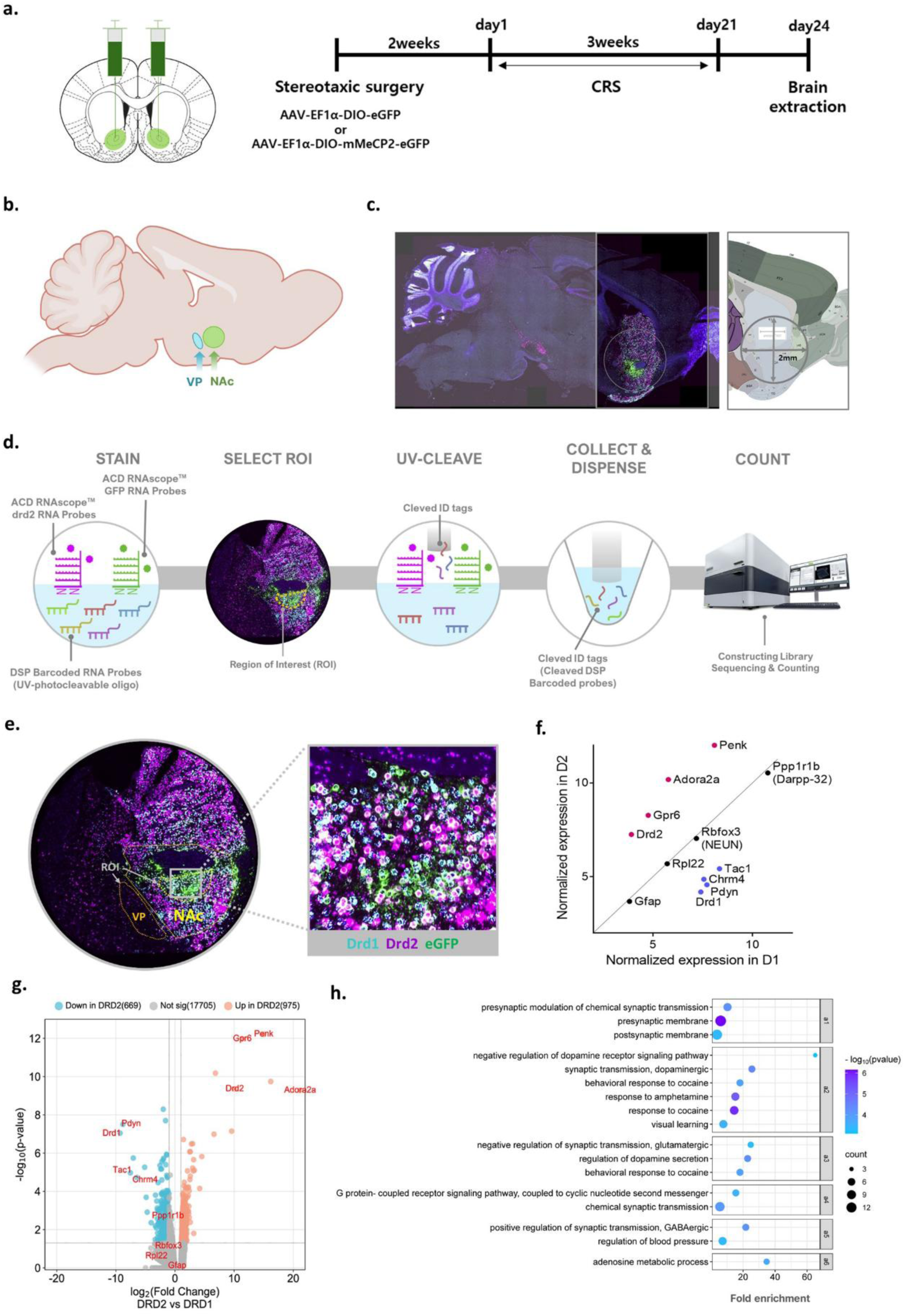
Procedure and validation for spatial transcriptome analysis of GeoMX DSP **a** Experimental procedure to confirm the effect of depression and genetic MeCP2 regulation. AAV expressing MeCP2 or eGFP was bilaterally injected into the NAc of D2R Cre mice using the Cre recombinase system. Brains were extracted 3 days after 3-week CRS exposure for the treatment groups (CC and CM) and prepared as formalin-fixed paraffin-embedded (FFPE) samples. **b** Experimental procedure for GEOMX analysis. RNA fluorescence in situ hybridization (FISH) technique was used to detect mRNA of genetic markers (drd2, drd1, nuclei) and marker for identifying AAV-infected cells (eGFP) and to distinguish target cells. mRNAs expressed in each cell were detected by their own UV-photoclevable DSP barcoded RNA probes. After setting a ROI in the NAc, UV was irradiated only for specific types of cells. UV irradiation detaches the barcode attached to the RNA probe, and sequencing analysis is performed by analyzing the read counts of the collected barcodes. **c** A sagittal section of the brain showing the target region for GEOMAX analysis. The transcriptome analysis included the NAc and VP regions (where the nerve terminals of D2R neurons project). **d** FISH images showing cells expressing D1R and D2R neurons and GFP in the brain. In the main experiment, only the target region was separated into a 2-mm-diameter circle as shown in the figure, and was used for analysis by GEOMX DSP. **e** Representative FISH images showing the ROI set for the target region (NAc, VP). For the group-wise analysis (NC-CC-CM) of D2R neurons in the NAc, transcriptome analysis was performed in cells coexpressing Drd2 and eGPF. **f, g** The expression levels of representative cell marker genes of each neuron were confirmed to verify the validity of Drd1 and Drd2 cells identified by GEOMX DSP technology in the NAc. **f** Graph showing the normalized expression (Log2) levels of representative genes for each cell type (D2R sample n = 9 mice, D1R sample n = 8 mice). **g** Volcano plot graph comparing gene expression levels of D2R neurons (left, blue). Genes with low expression levels (left, blue points) and high expression levels (right, red points) in D2R neurons compared to D1R neurons are indicated centered on the reference point (o) on the X-axis. The X-axis represents Log2 (fold change), and the Y-axis represents -log10 (p-value). **h** Results of DAVID functional annotation clustering analysis of 372 genes showing significant expression differences between D1R neurons and D2R neurons. [Fold change (FC)] > 1.5, p <0.05. a1-6: annotation clusters 1-6.

Before performing the DEG analysis on the CRS effect, we verified the validity of the cell type separation method using the GeoMX DSP method. We performed a comparative analysis of the expression levels of representative markers of D1R neurons and D2R neurons reported in previous studies.(Ho, Both *et al*., 2018; Kim, Wei *et al*., 2021). We confirmed that the distinguished cells closely reflected the molecular characteristics of each cell (Figs. 5f,g, Supplementary Figs. 12a,b). In the Log2 fold expression, Drd1 (9.29), Pdyn (8.82), Tac1 (7.55), Slc35d3 (7.06), Chrm4 (6.45), Foxp2 (4.70), and Sstr4 (3.30) were expressed at high levels in D1R neurons, and Adora2a (21.16), Penk (15.00), Gpr6 (11.38), Drd2 (10.10), and Grik3 (9.58) were expressed at high levels in D2R neurons. There was no difference in the expression of neuron (NeuN) and MSN (Darpp-32) specific genes between the two subtypes of cells, and a glia-specific marker, Gfap, showed a low expression level, revealing that the target cells were successfully distinguished.

We found 372 genes with significantly different expression levels between D1R and D2R neurons (300 upregulated genes, 72 downregulated genes FC>1.5, p≤0.05) (Fig. 5h, Supplementary Figs. 12c,d, Supplementary Table 1, 2). To identify the main functions of these genes, a DAVID functional annotation clustering analysis was performed on the GO database. Through this analysis, we clustered groups of genes with similar functions and ranked their functions based on the group enrichment score and the geometric mean (-log10 scale) of the p-values of the members of the corresponding annotation cluster. We confirmed that there were functional differences in synaptic modulation and transmission, addictive drugs, dopaminergic, glutamatergic, and GABAergic synaptic transmission, and G protein-coupled receptor signaling pathways between the two sub-cell types (p≤0.05, false discovery rate, FDR≤0.05) (Fig. 5h). Compared with D1R neurons, 300 genes upregulated in D2R neurons were enriched in the neuronal cell body, adenosin metabolic process, presynaptic and postsynaptic membranes, etc. (Supplementary Fig. 12c, Supplementary Table 3). In addition, 72 downregulated genes compared with D1R neurons were enriched in G-protein coupled receptor signaling, transcriptional regulation by RNA polymerase II, protein kinase activity, etc. (Supplementary Fig. 12d, Supplementary Table 4). There was no difference between D1R and D2R neurons in the mRNA expression level of MeCP2 (Supplementary Fig. 13a).

### MeCP2 regulates signaling and plasticity-related transcriptional networks in D2R neurons

To confirm the effects of CRS and rescuing overexpressed MeCP2 in D2R neurons, a DEG analysis was performed among three groups—normal control (NC), CRS-exposed (CC), and CRS-exposed with MeCP2 overexpression (CM)—specifically in eGFP-positive D2R neurons in the NAc. As expected, MeCP2 expression was significantly elevated in the CM group (p<0.0001), whereas no significant difference in MeCP2 mRNA levels was observed between the NC and CC groups. This indicates that the CRS-induced reduction in MeCP2 protein likely occurs via post-transcriptional regulation (Supplementary Fig. 13b).

In the DEG analysis we focused on the function of downstream targets regulated by MeCP2. We identified 457 genes ([FC]>1.3, p≤0.05; upregulated:325 genes, downregulated: 132 genes) altered by CRS in D2R neurons through the DEG analysis (Supplementary Fig. 14a, Supplementary Table 5). We also identified enriched cellular components associated with depressive symptoms using a GO analysis (Supplementary Fig. 14b, Supplementary Tables 6-8). The upregulated DEGs in D2R neurons by CRS were enriched for “ciliary base” and “ciliary basal body”, whereas the downregulated DEGs were enriched for “somatodendritic compartment”, “neuron to neuron synapse”, “postsynaptic density”, “asymmetric synapse”, and “synapse”. From the GO and UniProt databases (Universal Protein Resource), a DAVID functional annotation cluster analysis showed that CRS altered genes were related to ionotropic glutamate receptor binding, endoplasmic reticulum, proteasome complex, actin binding, palmitoylation, and peroxisome-related functions in D2R neurons (Supplementary Figs. 14c-e, Supplementary Tables 9-11).

Among 457 genes that showed different levels of expression between the normal (NC) and depressed (CC) groups, 250 genes (187 upregulated genes, 63 downregulated genes) showed attentuated effects of CRS by MeCP2 overexpression. This result suggests molecular-level restoration from depressive symptoms (CC vs. NC [FC]>1.3, p p≤0.05; CM vs. NC p>0.10) (Figs. 6a-d). The results of a heatmap analysis show that the expression levels of genes that were significantly increased or decreased in CC compared to NC were largely rescued in the CM group (Fig. 6a). We confirmed the molecular mechanism of MeCP2 underlying the alleviation of depressive symptoms through a functional analysis of 250 genes (Figs. 6b-e, Supplementary Table 12). The top five most enriched cellular components in the GO analysis were “neuron to neuron synapse”, “postsynaptic density”, “asymmetric synapse”, “postsynaptic specialization”, and “glutamatergic synapse” (Fig. 6b, Supplementary Tables 13-15). MeCP2 overexpression in D2R neurons showed the most dominant rescue effect on the synaptic function-related genes downregulated by CRS. In addition to synaptic function, a functional annotation cluster analysis was performed to summarize the rescued functions associated with MeCP2 upregulation (Figs. 6c, d, Supplementary Fig. 15a, Supplementary Tables 16-18).

**Fig. 6.**
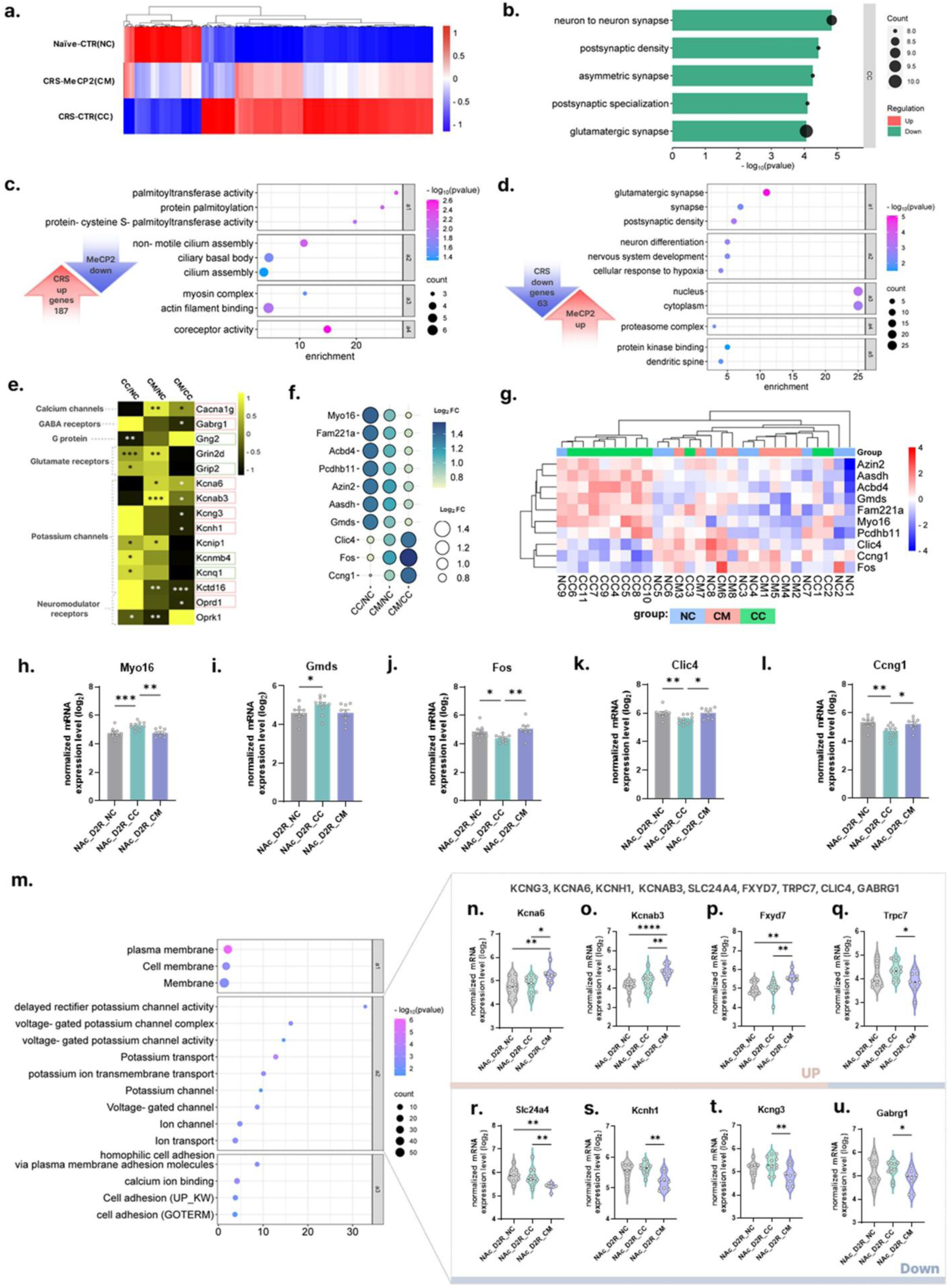
DEG analysis using GeoMX DSP. Comparison group: NC: Naive -CTR (control AAV), CC: CRS-CTR, CM: CRS-MeCP2 (AAV-MeCP2), n = 8, 11, 8 mice/group **a-d** 250 genes showing the rescue effect of MeCP2 in depression were selected. Genes whose expression levels were significantly changed by CRS (CC/NC, [FC]>1.3, p ≤0.05), but whose significance was lost after genetic MeCP2 increase (CM/NC, p>.10) were selected as analysis subjects. **a** Image showing the relative comparison of mRNA expression (Log2) between each group as a heat map. Heatmaps colored from blue to red according to Z-score scale −1 to 1. **b** Gene ontology enrichment analysis for cellular component in 250 genes[up(red) and down(green) genes by CRS exposure, no results in upregulated genes], False Discovery Rate (FDR) q <0.05 **c, d** Cluster analysis graph of DAVID functional annotation for 250 genes, X-axis represents enrichment scores for each cluster. Functional analysis was also performed on 187 up genes (c) and 63 down genes (d) in CC compared to NC **e** Heatmap image showing relative comparison of FC levels between groups of plasticity-related genes. Pink squared genes (CM vs. CC, p ≤0.05), green squared genes (CC vs. NC, p ≤0.05; CM vs. NC, p>0.10). Heatmaps colored from black to yellow according to Z-score scale −1 to 1 **f-m** Top 10 genes whose expression was restored by genetic MeCP2 increase (CC/NC [FC]>1.3, p ≤0.05, CM/NC p >0.30, CM/CC p ≤0.05). **f** 7 genes were downregulated and 3 were upregulated by MeCP2 upregulation. Bubble plots for Log2 FC values **g** Heatmaps for Log2 normalized expression values of individual NC-CC-CM samples. Heatmaps colored from blue to red according to Z-score scale −4 to 4. **h-l** Five genes whose expression was most significantly restored by genetic MeCP2 upregulation(CC/NC [FC] > 1.3, p ≤0.05, CM/NC p >0.40, CM/CC [FC] > 1.3, p ≤0.05), One-way ANOVA, post-hoc Fisher’s test, **h** Myo16: F(2, 25) = 10.06, p = 0.0006, NC vs. CC p = 0.0007, NC vs. CM p = 0.9612, CC vs. CM p = 0.0010, **i** Gmds: F(2, 25) = 2.956, p = 0.0704, NC vs. CC p = 0.0487, NC vs. CM p = 0.9676, CC vs. CM p = 0.0010, **j** Fos: F(2, 25) = 6.297, p = 0.0061, NC vs. CC p = 0.0253, NC vs. CM p = 0.3005, CC vs. CM p = 0.0513, **k** Clic4: F(2, 25) = 5.356, p = 0.0116, NC vs. CC p = 0.0099, NC vs. CM p = 0.9632, CC vs. CM p = 0.0109, **l** Ccng: F(2, 25) = 5.834, p = 0.0083, NC vs. CC p = 0.0036, NC vs. CM p = 0.5199, CC vs. CM p = 0.0226, All data are shown as mean ± SEM. **m-u** Analysis of the effect of genetic MeCP2 increase (CM) compared to CRS exposure group (CC). **m** DAVID functional annotation cluster analysis graph for 100 genes with altered expression in the CM group compared to the CC group (CM/CC, [FC]>1.3, p ≤0.05) **n-u** Comparison graph of normalized mRNA expression levels (Log2) of genes included in functional annotation cluster 2 (a2). One-way ANOVA, post-hoc Fisher’s test **n** Kcna6: F(2, 25) = 4.935, p = 0.0156, NC vs. CC p = 0.6784, NC vs. CM p = 0.0078, CC vs. CM p = 0.0147,**o** Kcnab3: F(2, 25) = 11.11, p = 0.0004, NC vs. CC p = 0.0896, NC vs. CM p <0.0001, CC vs. CM p = 0.0040 **p** Fxyd7: F(2, 25) = 6.985, p = 0.0039, NC vs. CC p = 0.2444, NC vs. CM p = 0.2104, CC vs. CM p = 0.0195 **q** Trpc7: F(2, 25) = 3.125, p = 0.0614, NC vs. CC p = 0.8515, NC vs. CM p = 0.0045, CC vs. CM p = 0.0020 **r** Slc24a4: F(2, 25) = 6.156, p = 0.0067, NC vs. CC p = 0.6325, NC vs. CM p = 0.0101, CC vs. CM p = 0.0146 **s** Kcnh1: F(2, 25) = 5.799, p = 0.0085, NC vs. CC p = 0.1818, NC vs. CM p = 0.0583, CC vs. CM p = 0.0022 **t** Kcng3: F(2, 25) = 4.432, p = 0.0225, NC vs. CC p = 0.2355, NC vs. CM p = 0.0975, CC vs. CM p = 0.0064 **u** Gabrg1: F(2, 25) = 2.294, p = 0.1217, NC vs. CC p = 0.5166, NC vs. CM p = 0.1678, CC vs. CM p = 0.0439. All data are shown as mean ± SEM. *p ≤0.05, **p <0.01, ***p <0.001, ****p <0.0001

**Fig. 7.**
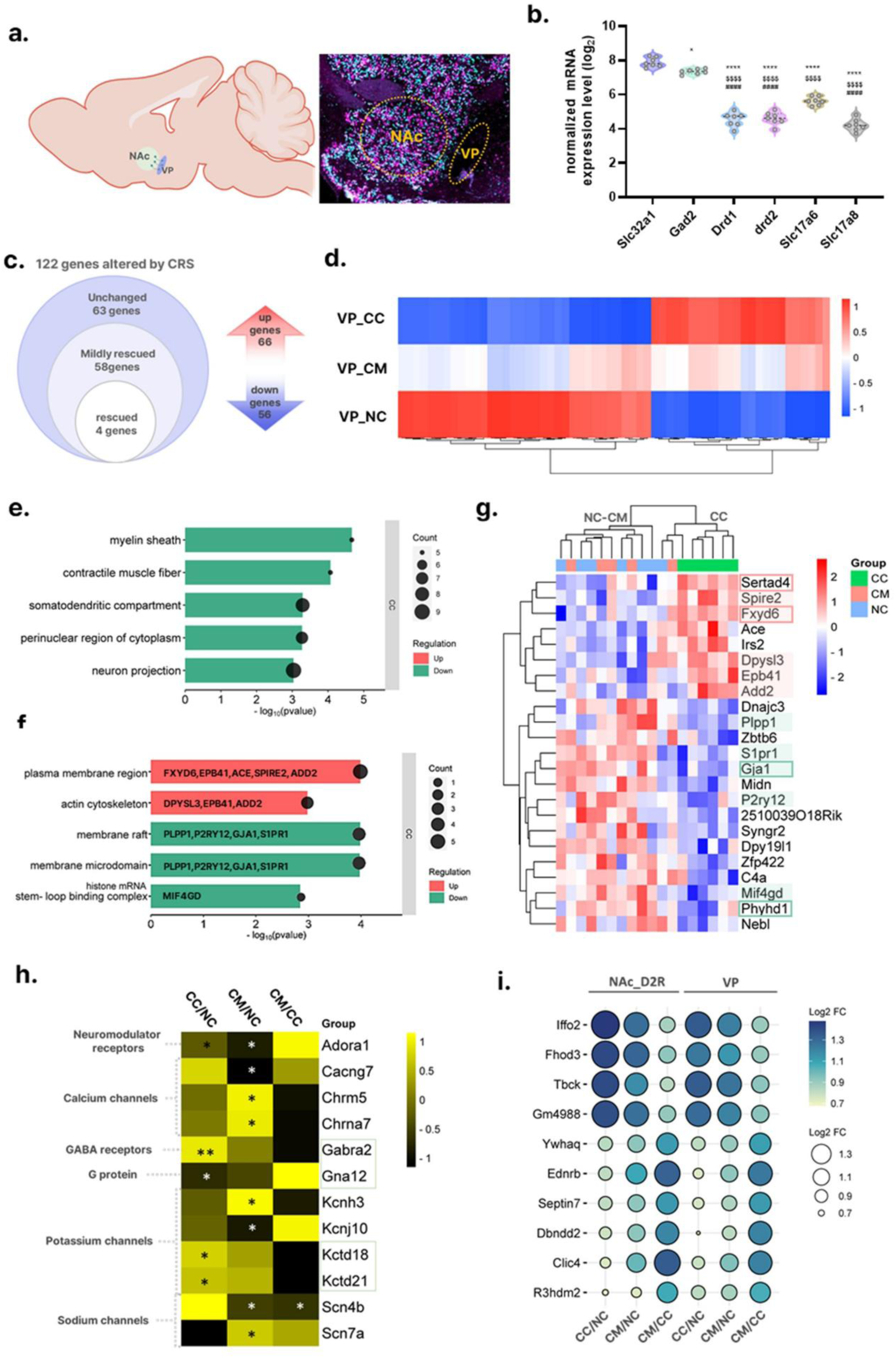
Molecular restoration of neural circuitry following MeCP2 overexpression. n = 7 (NC), 6 (CC), 5(CM) mice/group **a** Illustration showing the locations of the NAc and VP in sagittal sections (left) and representative FISH images indicating the ROI of the VP region to be sampled for GeoMX analysis (right panel) **b** Graph showing the expression levels of representative cell marker genes in VP. Samples from the NC group (n = 7) were used for the analysis. One-way ANOVA analysis, F(5, 36) = 171.8, p <0.0001, post-hoc Holm-Šídák’s test, Slc32a1 vs. Gad2 p = 0.0107, Slc32a1 vs. Drd1, Drd2, Slc17a6, Slc17a8 All have the same p value in each analysis. p ≤0.0001, Gad2 vs. Drd1, Drd2, Slc17a6, Slc17a8 All have the same p value in each analysis. p ≤0.0001, Drd1 vs. drd2 p = 0.7242, Drd1 vs. Slc17a6 p <0.0001, Drd1 vs. Slc17a8 p = 0.1172, drd2 vs. Slc17a6 p <0.0001, drd2 vs. Slc17a8 p = 0.0808, Slc17a6 vs. Slc17a8 p <0.0001, *p ≤0.05, ****p <0.0001 (compared to Slc32a1), $$$$p <0.0001 (compared to Gad2), ####p <0.001(compared to Slc17a6) **c** The expression levels of 122 genes (66 up genes and 56 down genes) of VP were significantly changed by CRS (CC vs. NC [FC]>1.3, p ≤0.05). Among them, 59 genes showed the alleviated CRS effect after genetic MeCP2 increase in D2R neurons of the NAc (CM sv. NC, p>0.10). **d** Heatmap images of 58 genes showing the alleviated CRS effect in VP after MeCP2 increase in NAc. Heatmaps colored from blue to red according to Z-score scale −1 to 1, **e** Gene ontology enrichment analysis for cellular component in 58 genes of VP. False Discovery Rate (FDR) q<0.05. Up (red bar, no results) and down (green bar) genes by CRS **f** Gene ontology enrichment analysis for cellular component in 23 genes of VP. False Discovery Rate (FDR) q<0.05. Up (red bar) and down (green bar) genes by CRS. Genes corresponding to each function are presented. **g** List of 23 genes whose expression levels were rescued by increased MeCP2 in the NAc. Log2 normalized expression values from individual samples were used (CC vs. NC [FC] > 1.3, p ≤0.05, CM vs. CC. 8 up and 15 down genes were presented. Pink (up) and green (down) shades are the list of genes derived from GO analysis (f). **h** Changes in plasticity-related genes in VP. Green boxes show genes with attenuated CRS effects (CC vs. NC. P ≤0.05, CM vs. NC *p ≤0.05, **p <0.01 (compared NC), Heatmaps colored from black to yellow according to Z-score scale −1 to 1. **i** Bubble plot graphs of 10 genes showing the same expression change pattern for depression and MeCP2 modulation effects in the NAc D2R neurons and VP (CC vs. NC, [FC]>1.2, p ≤0.05, CM vs. NC, p>.10, CM vs. CC, p ≤0.05).

In the functional annotation clustering analysis, among the genes upregulated by CRS, those whose expression was rescued by MeCP2 overexpression were enriched in the functional categories related to palmitoylation (group enrichment score: 1.815), cilium (1.530), and actin filament binding (1.112), in that order (Fig. 6c, Supplementary Table 17). Among the genes downregulated by CRS, the rescued gene set was enriched in clusters associated with glutamatergic synapse (3.527), nervous system development (2.543), nucleus (2.335), and proteasome complex (1.670), in that order (Fig. 6d, Supplementary Table 18). Recent studies have reported that palmitoylation (1.66) (Globa and Bamji, 2017; Sohn and Park, 2019), cillium (1.52) (Lucarelli, Di Pietro *et al*., 2019; Alhassen, Alhassen *et al*., 2023), and actin filament binding (1.112) (Globa and Bamji, 2017; Gentile, Carrizales *et al*., 2022) are related to the regulation of synaptic function. This suggests that restoring the expression of these genes may alleviate synaptic dysfunction.

The most representative neuropathological features of depression are synaptic dysfunction and abnormal neural activity(Duman and Aghajanian, 2012; Marsden, 2013). In our study, optogenetic activation of D2R neurons in the NAc revealed that mice exposed to chronic restraint stress (CC group) exhibited significantly reduced neural activity compared to controls (NC group), which was restored by MeCP2 overexpression (Fig. 4). To further elucidate the neurobiological role of MeCP2 in regulating neural activity and synaptic function, we conducted an extended analysis focused on these processes (Fig. 6e, Supplementary Figs. 16-18).

In the full functional annotation analysis of 250 genes, we estimated the protein-protein interaction (PPI) network for 22 genes related to glutamatergic synapses (eg, Rhoa, AKT1, Nr3c1, Dlg4, Il1rap) whose enrichment score was ranked the highest in the annotation group (Fig. 6d, Supplementary Fig. 15b, Supplementary Table 16). To understand the putative PPI, we identified a functional network from the STRING database. Genes were classified as members of networks related to axonal growth inhibition (RHOA activation), regulation of post-neurotransmitter receptor activity, and trans-synaptic signaling by trans-synaptic complexes. Alteration of genes constituting Rho family GTPases, including Rhoa, has been reported in recent studies using other depression models, suggesting that they are key factors mediating the regulation of depression(Fox, Chandra *et al*., 2020; Butto, Chongtham *et al*., 2024).

Next, we screened 236 plasticity-related genes encoding potassium, calcium or sodium channels and GABA, glutamate or neuromodulatory receptors or related G proteins(Câmara and Brandão, 2020), and derived 15 genes that were altered by CRS or modulated by MeCP2 (p ≤ 0.05) (Fig. 6e, Supplementary Figs. 16, 17, Supplementary Tables 19-25). CRS altered the expression of genes related to glutamate receptors (Grin2d, Grip2), potassuim channels (Kcnip1, Kcnmb4, Kcnq1), and neuromodulator receptors (Oprk1). In contrast, MeCP2 overexpression regulated genes constituting calcium channels (Cacna1g), GABA receptors (Gabrag1), potassuim channels (Kcna6, Kcnab3, Kcng3, Kcnh1, Kctd16), and neuromodulator receptors (Oprd1). These genes were not altered by CRS, and in particular, genes constituting subunits of potassium channels (8 out of 15 plasticity-related genes) were significantly affected by MeCP2 overexpression.

We derived 10 genes that were most resilient to CRS-induced degeneration due to MeCP2 overexpression (Figs. 6f-l). These genes satisfied three conditions: (i) they were significantly decreased by CRS (CC/NC, [FC]>1.3, p≤0.05), (ii) but their expression levels were not different from those of the control group (NC) due to MeCP2 overexpression (CM/NC, p>0.35), and (iii) at the same time, their expression levels were significantly restored to the control group level compared to the CRS group (CC) (CM/CC, p≤0.05). Among them, the expression levels of seven genes were decreased by MeCP2 overexpression (Myo16, Fam221a, acbd4, Pcdhb11, Azin2, Aasdh, Gmds), and three genes were increased (Clic4, Fos, Ccng1) (Figs. 6f, g). Among these, the final five genes with the highest expression changes in the MeCP2 overexpression group (CM) compared to the CRS group (CC) were Myo16, Gmds, Fos, Clic4, and Ccng1 ([FC] > 1.3, p≤0.05) (Figs. 6h-i). These genes are known to be mainly involved in cell survival and signal transduction. Myo16 is a signaling factor that integrates cell signaling pathways into actin cytoskeleton reorganization(Roesler, Lombino *et al*., 2019; Telek, Kengyel *et al*., 2020) and has been reported to be overexpressed in the brains of depressed suicidal patients(Gross, Pacis *et al*., 2017; Arčan, Kouter *et al*., 2022). Gmds is the first enzyme that converts GDP-Mannose to GDP-4-Dehydro-6-Deoxy-Mannose. It also affects the glycosylation process and has been reported to be involved in cell proliferation and survival(Wei, Zhang *et al*., 2018). Fos showed a decrease in expression in the CRS group (CC) and a recovery in expression in the MeCP2 overexpression group. CM immediate early genes (IEGs) such as Fos are representative genes reflecting the level of neural activity in the brain. The gene expression changes observed across the three groups (NC, CC, and CM) are consistent with electrophysiological findings (Fig. 4). This supports the reliability of the results showing the changes in neural activity and their direction of change due to CRS exposure or MeCP2 overexpression. Among the differently expressed genes, Clic4 (chloride intracellular channel 4) encodes a chloride ion channel that, although not fully characterized, has been implicated in neuronal function, including intracellular oxidative stress responses, inflammatory signaling pathways, and calcium signal modulation(Shukla and Yuspa, 2010; Xue, Lu *et al*., 2016). Ccng1 (Cyclin G1), a cell cycle regulatory protein, has also been reported to regulate DNA damage during the cell cycle and to be involved in inflammatory responses (Horne, Goolsby *et al*., 1996; Ohtsuka, Jensen *et al*., 2004).

To confirm the direct effect of MeCP2 overexpression upon CRS exposure, we performed a DEG analysis between the CRS group (CC) and the group with combined CRS and MeCP2 overexpression (CM) (CM/CC, [FC]>1.3, p≤0.05) (Figs. 6m-u, Supplementary Table 16). We derived 100 DEGs whose expression was significantly changed by MeCP2 overexpression, and performed a functional analysis on these genes (Figs. 6m-u, Supplementary Fig. 26). In the GO analysis, the enriched functions in the CM group compared to the CC were “synapse assembly”, “cell junction assembly”, and “potassium ion transport” in biological processing, and “synapse”, “postsynaptic membrane”, “synaptic membrane”, “glutamatergic synapse”, and “postsynaptic density membrane” in the cellular components, in that order (Supplementary Fig. 18b). These functions were predominantly expressed in DEGs downregulated by MeCP2 overexpression (Supplementary Fig. 18c). In the functional annotation cluster analysis, the notable functional changes in the MeCP2 overexpression group (CM) compared to the CRS group (CC) were related to membranes (enrichment score for annotation cluster: 3.840), potassium and ion channels or transporters (2.427), and calcium ion binding and cell adhesion (2.380) (Fig. 6m). In particular, MeCP2 overexpression in the CM compared to the CC mainly regulated the voltage-gated potassium (K⁺) channel-related genes, Kcng3, Kcnh1, Kcna6, and Kcnab3 (Figs. 6n,o,s,t). In addition, genes related to signal transduction of ion channels and maintenance of homeostasis, Slc24a4, a Na⁺/K⁺/Ca²⁺ exchanger, Trpc7, a member of the TRP channel family that mediates calcium influx, Fxyd7, a modulator of Na⁺/K⁺-ATPase, and Gabrg1, the primary inhibitory neurotransmitter receptor, were regulated by the increase in MeCP2 (Figs. 6p-r,u).

The functional annotation group for upregulated genes in CM compared to CC consisted of DEGs related to potassium and ion transport (group enrichment score: 1.183 (Supplementary Fig. 18d), and downregulated genes were enriched in cell adhesion (2.838), axon guidance (1.265), and potassium transport (1.175) (Supplementary Fig. 18e).

When we integrated the results of several DEG analyses that we performed to explore the rescue effect of MeCP2 in CRS, MeCP2 overexpression in D2R neurons of the NAc in the CRS model showed that DEGs related to synaptic function were most enriched. Specifically, MeCP2 overexpression alleviated the CRS effect of DEGs that were downregulated by CRS. This included the regulation of glutamatergic synapses and potassium-mediated signaling systems as important neurobiological rescue mechanisms. In addition, MeCP2 overexpression alleviated CRS effects in DEGs associated with palmitoylation, cilia, and actin filament binding. In summary, in D2R neurons of the NAc, MeCP2 is involved in the regulation of synaptic function, cell structure, and survival pathways altered by CRS.

### MeCP2 overexpression in D2R neurons in the NAc promotes molecular rescue at the circuit level in depression

Our data demonstrate that MeCP2 overexpression in D2R neurons of the NAc normalizes the behavioral phenotype of CRS-induced depressive symptoms and restores the expression of related genes in various functions, including synapses. As mentioned above, we investigated whether functional recovery of one region can lead to circuit-level recovery in terms of molecular recovery. The ventral pellidum (VP) is a direct projection area of D2R neurons in the NAc and has recently been reported to be associated with depression (Fig. 7a) (Knowland, Lilascharoen *et al*., 2017; Morais-Silva, Campbell *et al*., 2023).

To validate the reliability of the established ROI, we first checked the expression levels of representative marker genes of VP cells by referring to recent literature(Ottenheimer, Simon *et al*., 2024; Yang, Fang *et al*., 2024). The VP is composed of approximately 90% GABAergic neurons and 10% glutamatergic neurons. Accordingly, we compared the expression levels of canonical GABA-related marker genes, such as Gad2 (involved in GABA synthesis) and Slc32a1 (encoding the vesicular GABA transporter), as well as glutamate-related marker genes Slc17a6 and Slc17a8 (encoding vesicular glutamate transporters, VGLUT2 and VGLT3), with those of dopamine receptor genes Drd1 and Drd2 (Fig. 7b). Similar to previous reports, the expression levels of Gad2 and Slc32a1 were significantly higher than those of Slc17a6, Slc17a8, Drd1, and Drd2 (p<0.0001). The expression level of Slc17a6 was significantly higher than those of Drd1 and Drd2 (p<0.0001). These results show that the sampled tissues accurately reflect the cellular composition of the VP.

From the results of the DEG analysis, we confirmed that the expression of 122 genes was altered by CRS in VP (upregulated genes: 66, downregulated genes: 56, [FC]>1.3, p≤0.05) (Fig. 7c, Supplementary Table 27). In the GO analysis, these DEGs were mainly altered in terms of myelination and ribosome-related functions, such as “myelin sheath”, “cytosolic ribosome”, “cytosolic small ribosomal subunit”, “ribosomal subunit”, and “melanosome” in the cellular components (Supplementary Fig. 19a, Supplementary Tables 28-30). In the results of the functional annotation clustering analysis, the main cellular and biological functions of the genes expressed in the VP were altered, such as the ribosome (enrichment score for annotation group1: 2.19) and cell body (1.14). Furthermore, the PPI analysis for annotation group 1 confirmed a decrease in the expression of genes in the ribosome-related network (Supplementary Fig. 19b, Supplementary Table 31). Among 122 DEGs, a total of 58 DEGs were attenuated by the CRS effect in VP due to MeCP2 overexpression in the NAc (CM/NC, p>.10) (Figs. 7c,d). In the GO analysis of these genes, DEGs in the cellular component were enriched for CRS in the “myelin sheath”, “contractile muscle fiber”, “somatodendritic compartment”, “perinuclear region of cytoplasm”, and “neuron projection” (Fig. 7e, Supplementary Tables 32-35).

Among the genes altered in CC compared to NC, 23 genes were significantly rescued in CM compared to CC, and they showed no difference in expression levels compared to NC (CC vs. NC [FC]>1.2, p≤0.05; CM vs. NC, p>0.10; CM vs. CC p≤0.05) (Figs. 7f, g, Supplementary Table 36). In the GO analysis, upregulated DEGs were enriched in the “plasma membrane region” and “actin cytoskeleton” in cellular components, and downregulated DEGs were enriched in the “membrain raft”, “membrane micordomain”, etc. The top four genes that showed the most pronounced rescued effects upon MeCP2 overexpression in the NAc were Sertad4 (SERTA Domain Containing 4)(Chen, Zheng *et al*., 2024), which is involved in immune response, Phyhd1 (Phytanoyl-CoA 2-Hydroxylase Domain Containing 1) encoding dioxygenase involved in fatty acid metabolism and in immune response regulation, Fxyd6 (FXYD domain containing ion transport regulator 6), which acts as a regulator of Na+/K+-ATPase complex and is involved in maintaining intracellular ion balance, and Gja1 (gap junction protein alpha 1), which is involved in the formation of gap junctions essential for intercellular signaling ((Fig. 7g, Supplementary Figs. 20a-e).

We confirmed that MeCP2 overexpression in D2R neurons of the NAc regulated several genes related to synaptic plasticity in the VP (e.g. Gabra2, Gna12, Kctd18, Kctd21). This suggests that synaptic remodeling occurred at the circuit level (Fig. 7h). Given that D2R neurons in the NAc are GABAergic neurons, the changes in Gabra2 expression in the VP may reflect a direct neuroplastic response to changes in neuronal activity in the NAc induced by CRS.

We also searched for genes that showed the same degeneration and restructuring patterns in response to CRS and MeCP2 overexpression in the NAc and VP (Fig. 7i, Supplementary Figs. 20f-h). Ten genes showed similar patterns of alteration and rescue in both regions (CC vs. NC, [FC]>1.2, p≤0.05, CM vs. NC, p>.10) (Fig. 7i, Supplementary Table 37). In addition, among the top 10 representative genes restored by MeCP2 overexpression in D2R neurons in the NAc (Figs. 6f,g), only Clic4 in the VP showed a similar pattern of increase and decrease (CC vs. NC FC=1.2731, p=0.0168, CM vs. NC p=0.7500) (Supplementary Figs. 20f-h).

In summary, MeCP2 overexpression in D2R neurons in the NAc restored a significant number of genes whose expression was altered by CRS in the adjacent region, the VP. This indicates that MeCP2 regulation in the NAc induced molecular recovery not only in the NAc but also in directly connected neural circuit regions. CRS altered the expression of genes involved in myelination and ribosome and synaptic functions in the VP, whereas MeCP2 in the NAc restored the expression of a significant number of genes responsible for these functions.

## Discussion

In this study, we demonstrated MeCP2 has a critical role in regulating depressive-like behaviors induced by chronic restraint stress (CRS) in the nucleus accumbens (NAc). CRS exposure led to depressive-like phenotypes—such as weight loss, increased anxiety-like behavior, and behavioral despair. Importantly, MeCP2 protein expression was selectively reduced in D2R-expressing neurons of the NAc. Bidirectional genetic manipulation of MeCP2 in these neurons confirmed its causal role: MeCP2 knockdown alone induced depressive-like behaviors whereas MeCP2 overexpression rescued the CRS-induced behavioral phenotype. These results position MeCP2 as a key regulator of depressive behaviors in D2R neurons of the NAc.

We next sought to determine the neurophysiological basis for this behavioral rescue. Using optogenetic activation and multielectrode array recordings, we found that CRS reduced the activity of D2R neurons in the NAc, which was restored by MeCP2 overexpression. These findings align with previous studies showing that MeCP2 deficiency reduces excitatory synaptic transmission and plasticity, while MeCP2 overexpression enhances these functions in other brain regions(Na, Nelson *et al*., 2013; Huang, Zhang et al., 2021). For instance, MeCP2 deficiency has been associated with decreased spontaneous excitatory postsynaptic current (mEPSC) frequency and impaired plasticity in hippocampal neurons(Nelson, Kavalali *et al*., 2006), while MeCP2 overexpression enhanced synaptic plasticity and cognitive performance in aging models(Huang, Zhang et al., 2021). However, MeCP2 has also been reported to have divergent effects depending on brain region and cell type(Chao, Zoghbi *et al*., 2007; Kline, Ogier *et al*., 2010; Wood and Shepherd, 2010; Zhang, Zak *et al*., 2010; bSugino, Hempel *et al*., 2014; Calfa, Li *et al*., 2015). For example, MeCP2 deletion decreases excitability in hippocampal CA1 neurons and cortical pyramidal cells(Chao, Zoghbi et al., 2007; Wood and Shepherd, 2010), but increases excitability in the CA3 region and the nucleus of the solitary tract(Kline, Ogier et al., 2010; Calfa, Li et al., 2015). Additionally, MeCP2 loss reduces GABAergic inhibition in the thalamus(Zhang, Zak et al., 2010). Our prior work also demonstrated that pan-neuronal MeCP2 knockdown in the dorsal striatum increases overall activity levels(Lee, Kim *et al*., 2022). Taken together, these findings highlight that MeCP2 plays a dynamic and context-dependent role in regulating neuronal excitability, reinforcing the importance of studying its function in a brain-region and cell-type-specific manner.

To elucidate the molecular basis of MeCP2’s rescue effects, we conducted spatial transcriptomic profiling of D2R neurons from normal (NC), CRS-exposed (CC), and rescued (CM) mice. MeCP2 overexpression reversed the expression of numerous genes that were dysregulated by CRS, particularly those related to synaptic function. Among the most significantly restored were genes involved in glutamatergic synaptic transmission and potassium channel activity. Given that glutamatergic signaling is a well-established target for antidepressant interventions(Kendell, Krystal *et al*., 2005; Mathews, Henter *et al*., 2012) , our findings underscore the role of MeCP2 in modulating glutamatergic signaling pathways implicated in the pathophysiology of depression.

Future studies to clarify how MeCP2 influences glutamatergic transmission and how this contributes to the regulation of depressive-like behavior are warranted. Another key finding was that potassium channel-related gene expression changes between the CC and CM groups were strongly enriched. Potassium channels have recently emerged as promising therapeutic targets for depression (Tan, Costi *et al*., 2020; Zhang, Zhu *et al*., 2024). In our study, MeCP2 manipulation led to both up- and downregulation of specific potassium channel-related genes, suggesting a nuanced role in ion channel homeostasis. These results point to a broader functional relevance of MeCP2 in modulating neuronal excitability and synaptic plasticity and call for further investigation of its role in potassium signaling pathways within depression-related circuits.

In addition, MeCP2 overexpression attenuated CRS-induced increases in gene categories associated with palmitoylation (1.66)(Globa and Bamji, 2017; Buszka, Pytyś *et al*., 2023), cilium structure and function (1.52) (Lucarelli, Di Pietro et al., 2019; Alhassen, Alhassen et al., 2023), and actin filament binding (1.112) (Globa and Bamji, 2017; Gentile, Carrizales et al., 2022) —biological processes known to influence synaptic plasticity. These findings suggest that MeCP2 may alleviate CRS-induced reductions in neural activity by modulating a coordinated network of cellular pathways involved in synaptic regulation, ultimately contributing to the reversal of depressive-like behaviors. Among the most notable findings of this study is that MeCP2 overexpression in D2R neurons of the NAc effectively rescued depressive symptoms by regulating the expression of multiple synaptic function-related genes. Supporting this, transcriptomic analysis revealed that the expression of Fos— a well-established marker of neuronal activity—was downregulated in the CC group but restored to control levels in the CM group. These molecular changes are consistent with electrophysiological recordings, reinforcing the conclusion that MeCP2 overexpression counteracts CRS-induced deficits in neuronal activity within this cell population.

In an extended analysis, we investigated whether functional recovery in one brain region could induce circuit-level molecular changes. Genetic overexpression of MeCP2 in the NAc not only normalized CRS-induced depressive-like behaviors to control levels but also restored the expression of numerous genes altered by stress in the NAc. Based on the understanding that behavioral outcomes emerge from coordinated activity across interconnected regions, we hypothesized that rescuing molecular changes in the NAc could also lead to recovery in downstream circuits. To test this, we examined the ventral pallidum (VP)—a major projection target of D2R neurons in the NAc—which has recently been implicated in mood regulation and motivational processing(Morais-Silva, Campbell et al., 2023; Soares-Cunha and Heinsbroek, 2023). Our spatial transcriptomic analysis revealed that CRS induced significant alterations in myelination, ribosomal activity, and synaptic function-related genes in the VP, many of which were normalized in the CM group. These findings suggest that MeCP2-mediated rescue in the NAc can propagate circuit-level molecular recovery. Although our current findings provide evidence of molecular rescue beyond the NAc, a more comprehensive investigation is needed. Evaluating the circuit-wide consequences of target gene modulation will be essential for fully understanding therapeutic mechanisms and for guiding future intervention strategies. Spatial transcriptomics, as demonstrated here, offers a powerful tool to map these effects. Future studies should aim to characterize transcriptomic alterations in additional regions that are functionally connected to the NAc—such as the medial prefrontal cortex (mPFC), ventral tegmental area (VTA), and hippocampus—which are known to play critical roles in depressive symptomatology and motivational regulation.

Interestingly, while MeCP2 protein was significantly reduced in D2R neurons following CRS, mRNA levels were unchanged, suggesting a post-transcriptional mechanism. Previous studies have implicated miRNAs such as miR-132 and miR-124 in MeCP2 regulation, both of which are upregulated in the depressed brain. (Yang, Liu *et al*., 2020; Tong, Li *et al*., 2021). Exploring these miRNA-mediated mechanisms may further clarify upstream regulatory processes contributing to MeCP2 dysfunction in stress-related disorders.

This study is the first to demonstrate the therapeutic effect of MeCP2 modulation in a depression model, and to map its transcriptomic and electrophysiological consequences in a cell-type-specific manner. Future studies should investigate how MeCP2 modulates synaptic function in the NAc under chronic stress conditions, utilizing various electrophysiological techniques. Additionally, functional validation of individual MeCP2 target genes identified by transcriptomic analysis is necessary, and epigenetic approaches will be critical for these efforts. Since synaptic degeneration is a hallmark of depression, understanding how MeCP2 modulates synaptic integrity in specific neural circuits may advance both mechanistic understanding and therapeutic development. Collectively, our findings highlight MeCP2 as a central molecular switch regulating depression-like symptoms in D2R neurons of the NAc, with implications for developing cell-type-specific interventions.

## Acknowledgements

This work was supported by the Ministry of Food and Drug Safety (grant no. 25212MFDS003), the National Research Foundation of Korea (grant nos. RS-2024-00332024 and RS-2024-00463082), and the Korea Institute of Science and Technology (grant no. 2E33702).

We thank Eonji Noh and Seungyoun Lee of Theragen Bio for their assistance with bioinformatics analysis.

Brain diagrams were adapted from *The Mouse Brain in Stereotaxic Coordinates* (Paxinos and Franklin) and the Allen Brain Atlas (https://atlas.brain-map.org/).

Some figures were created using **BioRender.com.**

## Author information

### Authors and Affiliations

**Center for Brain Disorders, Brain Science Institute, Korea Institute of Science and Technology, Seoul, South Korea**

Jinhee Bae, Sun A Jung, Nazarii Frankiv, Eun Mi Hwang, Sangjoon Lee & Heh-In Im

**Biomaterials Research Center, Korea Institute of Science and Technology, Seoul, South Korea**

Sung Hoon Kim & Young-Min Kim

**Division of Bio-Medical Science & Technology, KIST School, Korea University of Science and Technology, Seoul, South Korea**

Sun A Jung, Nazarii Frankiv, Eun Mi Hwang, Young-Min Kim, Sangjoon Lee & Heh-In Im

### Contributions

J.B. contributed to study design, behavioral experimentation, data collection and analysis, data interpretation, and manuscript writing and revision. S.H.K. contributed to data collection and analysis, data interpretation, and manuscript revision. S.A.J. and N.F. contributed to data collection and analysis. E.M.W. contributed to AAV vector design and production, and manuscript revision. S.J.L. contributed to data analysis and interpretation, and manuscript revision. H.I.I. contributed to study design, data analysis and interpretation, and manuscript writing and revision. All authors reviewed and agreed to the final version of the manuscript.

### Corresponding author

Correspondence to Heh-In Im

## Ethics declarations

### Competing interests

The authors declare the following patent applications related to the findings of this study, all filed by the Korea Institute of Science and Technology:

- Application nos. 10-2023-0055806 (Republic of Korea) and 18/598,746 (United States), covering the use of MECP2 as a biomarker for depression diagnosis and as a therapeutic target.
- Application no. 10-2025-0003282 (Republic of Korea), covering the development of a Cre-dependent viral system for gene regulation of MeCP2; a corresponding PCT/US application is in progress.

### Data availability

Processed results of the transcriptomic analysis are available in the Supplementary Data files. Due to the size and nature of the raw sequencing data, they are available from the corresponding author upon reasonable request.

