## Supplemental Data 1 for "MeCP2 regulates cell type-specific functions of depressive-like symptoms in the nucleus accumbens"

Bae et al.

This file includes:

Material and Methods

SI Reference

Supplementary Figures. 1-20

### Material and Methods

A full description of all experimental procedures is provided in the Supplementary Information.

#### Animals

Seven to nine week old male C57BL/6J (DBL, Gyeonggi, South Korea), B6.FVB(Cg)-Tg(Drd2-cre)ER44Gsat/Mmcd(# 032108-UCD, MMRRC, CA, USA) or B6. Cg-Tg(Drd1-cre)262Gsat/Mmcd(# 030989-UCD, MMRRC, CA, USA)) mice were used for experimental procedures. All transgenic mice for experiments were backcrossed to wild-type C57BL/6J mice for several generations. All mice were housed under standard conditions at 21-23℃, 50-55% humidity, and a 12-hour light/dark cycle. All behavioral procedures were performed during the dark cycle. All experimental procedures were approved by the Animal Care and Use Committee of the Brain Science Institute of the Korea Institute of Science and Technology (KIST).

#### Chronic restraint stress model

Mice were restrained in a square restraint device with an internal area of 3 (width) × 3 (length) × 17 (length). They were secured inside the restraint device by a movable fixation wall of approximately 4.0 cm so that they could not move freely. In the space inside the restraint device, the mice could slightly change their body position but could not move forward or backward. The mice were held simultaneously for 6 h per day for 21 days. The control group received no disturbance except for 3 min of handling per day during the same period. Both groups received regular cage maintenance and body weight measurements during the CRS exposure period.

#### Behavioral Tests

##### Open Field Test

The locomotor activity of mice was analyzed using an open field test. The behavioral box of the open field test consisted of a white acrylic square arena (40 cm X 40 cm) and a 40 cm white wall. On the day after the CRS session, both the CRS and CTR groups were allowed to freely explore the box for 30 min, and their movements were recorded with an infrared camera. The analysis was performed using EthoVision software (Noldus, Wageningen, NL). The center zone was assigned to the central square area (20 cm X 20 cm). Locomotor activity was measured as the total distance traveled and the distance traveled in the center zone as dependent variables.

**Elevated Plus Maze Test**

An elevated plus maze (EPM) test was conducted to determine the anxiety level of mice. The EPM box consisted of two white open arms and two black closed arms. The closed arms had a 15 cm high wall and the open arms had no wall. The height of the box was 44 cm. The mice were placed in the central area of the EPM box, and the movements of the mice were recorded for 10 minutes using a video camera. The time spent in the open and closed arms was measured manually, and time in the arms was only recorded when all four paws of the mice entered the arms. The time spent in the central area was excluded from the measurement.

**Forced Swim Test**

A forced swim test (FST) to measure depressive symptoms in mice was performed in a cylinder 20 cm high and 15 cm in diameter for six minutes. The water temperature was 24±1°C, and fresh water was replaced immediately after each trial. Immobility was recorded during the last four minutes of the 6-minute period and analyzed manually. Mobile time was measured when the animals swam or moved by moving at least three limbs, including both forelimbs.

**Novel Object Recognition Test (NOR)**

A novel object recognition test was used to measure cognitive function. In the training trial (familiarization phase), mice were allowed to explore two identical objects placed at regular intervals in a white square box (40 × 40 × 40 cm, center: ~5 lx; periphery: ~4 lx) for 10 min. The test trial (recognition phase) was performed 24 h after the training trial for 10 min. Mice explored two different objects (familiar and novel objects). The behavior of mice was video recorded, and the smell time of the familiar or novel object was manually analyzed and compared during the test phase.

**Stereotaxic injection and adeno-associated viruses**

Under ketamine-xylazine anesthesia, stereotaxic injections of virus into the NAc were performed using a stereotaxic device (David Kopf Instruments). First, 500 nl of concentrated virus solution was injected bilaterally into the NAc (AP +1.4 mm; ML ±1.15 mm; VP -4.1 mm) using a syringe pump (Harvard Apparatus) at a slow rate (100 nl/min). The needles were slowly withdrawn 10 min after the injection. In experiments requiring insertion of an optogenetic cannula, the needles were removed and the optogenetic cannula was implanted bilaterally into the NAc (AP +1.4 mm; ML ±1.15 mm; VP -3.8 mm). Mice were used two to three weeks after AAV injection. We constructed a Cre-dependent AAV vector that we previously developed by inserting a scrambled or mouse MeCP2 shRNA sequence (5'GTCAGAAGACCAGGATCTC-3') for knockout of MeCP2 (AAV-GFP-creon-mMECP2 shRNA) (Zhou, Hong *et al.*, 2006; Kim, Kim *et al.*, 2022). To increase the expression of MeCP2, the full-length sequence of mouse MECP2 was inserted into the pAAV-Ef1a-DIO-EGFP-WPRE-pA (Plasmid #37084, Addgene, MA, USA) vector. All vectors were packaged as serotype DJ at the KIST Virus Facility (http://virus.kist.re.kr).

**Western Blot**

To analyze proteins in each sample, 40 μg of protein was added to 5X SDS-PAGE loading buffer and loaded onto 12% Tris-glycine SDS-polyacrylamide gel. The gel was transferred to an immibilion-P transfer membrane (IPVH 00010, Millipore), and the membrane was blocked with 5% SKIM milk in 0.03% Tris-Bisol-Tris (Tween 20, P2287, Sigma). The membrane was incubated overnight at 4°C with shaking in blocking buffer containing primary antibodies (rabbit anti-MeCP2 polyclonal antibody, 1:500, 07-013, Millipore; mouse anti-GAPDH, 1:5000, sc-47724, Santa Cruz, TX, USA; mouse anti-β actin, 1:5000, sc-47778, Santa Cruz, TX, USA). Membranes were washed three times in TBS-T with shaking for 10 min each and then incubated with secondary antibodies (goat anti-rabbit IgG-HRP, 1:5000, ab97051, Abcam, Cambridge, UK or goat anti-mouse IgG-HRP, 1:5000, sc-2005, Santa Cruz, TX, USA) with shaking at room temperature for 2 h. Membranes were rinsed again and visualized using SuperSignal West Pico Chemiluminescent Substrate (Thermo Scientific, MA, USA) according to the manufacturer's instructions. Immunoblots were detected with a luminescent image analyzer ImageQuant LAS4000 (GE Healthcare Life Sciences, BUX, UK). Densitometric quantification was performed and blots were analyzed with Image J software.

**Formalin-fixed paraffin-embedded tissue preparation**

Mouse brains were collected and fixed in 4% paraformaldehyde (PFA) for 24 hours at 4 °C. For paraffin embedding, brains were sequentially dehydrated using 40% ethanol, 70% ethanol, and a second 70% ethanol wash, each for 24 hours. Whole brains were then transferred to embedding cassettes and processed using a Tissue-Tek VIP Tissue Processor (Sakura Finetek). The 12-hour processing cycle included one 30-minute ethanol wash, five additional 1-hour ethanol incubations, three 45-minute CitriSolv washes (Decon Labs), and three 1-hour paraffin wax infiltrations. For GeoMx DSP, tissues were paraffin-embedded, and 2-mm-diameter circles of thalamic brain regions, including the NAc and VP, were punched out from individual paraffin blocks and embedded into tissue microarray (TMA) blocks to facilitate simultaneous processing and analysis.

**Immunohistochemistry**

Mice were deeply anesthetized with Avertin (2, 2, 2-tribromoethanol, Sigma) 2 h or three days after the CRS session. Mice were transcranially perfused with ice-cold 1x phosphate-buffered saline

(PBS) and then fixed in ice-cold 10% formalin solution (Sigma). Brains were postfixed overnight in 10% formalin solution at 4°C and stored in 30% sucrose. They were then sectioned (40 μm) using a microtome (Leica) at -20°C. Brain sections were incubated overnight at 4°C with primary antibodies (rabbit anti-MeCP2, 1:250, Millipore, 07-013; mouse anti-MeCP2, 1:250, Abcam, ab50005; rabbit anti-D1DR, 1:50, Abcam, ab20066; rabbit anti-D2DR, 1:50, Santa Cruz, sc-5303; chicken anti-GFP, Abcam, ab13970) and then probed with secondary antibodies (Alexa Fluor 488 goat anti-rabbit IgG, 1:400, Life Technologies, A-11008; Alexa Fluor 594 goat anti-rabbit IgG, 1:400, Life Technologies, A-11012; Alexa Fluor 488 goat anti-mouse IgG, 1:400, Life Technologies, A-11034; Alexa Fluor 594 goat anti-mouse IgG, 1:400, Life Technologies, A-11005; goat-anti-chicken IgG, 1:400, Abcam, ab150169) for 2 h at room temperature. Sections were mounted on coverslips using a mounting medium containing DAPI solution (H-1500, Vector). Confocal images were taken using a Zeiss LSM800 confocal microscope. For paraffin-embedded tissues, sections were deparaffinized in xylene and ethanol (100%, 95%, 70%, and 50%, respectively), and immunostaining was performed in the same manner as above for the brain tissue fixed on the slices.

**RT-qPCR**

Total RNA was extracted from homogenized brain tissues using TRIzol reagent (Life Technologies, NY, USA), according to the manufacturer's instructions. Complementary DNA (cDNA) was synthesized from the extracted RNA using the ReverTra Ace™ qPCR RT Master Mix (TOYOBO, Osaka, Japan). Quantitative real-time PCR (qPCR) was performed using THUNDERBIRD™ SYBR® qPCR Master Mix (TOYOBO) on a CFX Connect™ Real-Time PCR Detection System (Bio-Rad, CA, USA). The following primer pairs were used: MeCP2 (forward: 5′-GAGAGAGCAGAAACCACCTA-3′; reverse: 5′-TCTGATGCTGCTGCCTTT-3′), Drd2 (forward: 5′-ACTTGTGTGCCATCAGCATC-3′; reverse: 5′-AAGGACAGGACCCAGACGAT-3), Gapdh (forward: 5′-GACATCAAGAAGGTGGTGAAGC-3′; reverse: 5′-ACCACCCTGTTGCTGTAGCC-

3′). Relative mRNA expression levels were calculated using the ΔΔCt method, with Gapdh serving as the internal reference gene.

**FACs cell sorting**

Mice genetically engineered to express fluorescent reporter proteins under a cell type-specific promoter were used for fluorescence-based cell isolation. Animals were euthanized in accordance with institutional animal care guidelines, and brain tissues were harvested immediately post-mortem under sterile conditions. Tissues were mechanically dissociated using fine scissors or razor blades, followed by enzymatic digestion in RPMI 1640 or DMEM supplemented with 1–2 mg/mL collagenase D and 100 μg/mL DNase I at 37 °C for 30–45 minutes with gentle agitation. The resulting cell suspensions were filtered through a 70 μm cell strainer to obtain single-cell suspensions and centrifuged at 300–400 × g for five minutes. When required, erythrocyte lysis was performed using ACK lysis buffer for 1–3 minutes at room temperature. Cells were washed and resuspended in FACS buffer (PBS containing 2% FBS and 2 mM EDTA) and then passed through a 35 μm mesh to remove aggregates prior to sorting. Fluorescence-based cell sorting was performed on a MA900 Cell Sorter (SONY Biotechnology, Japan) equipped with appropriate lasers and filters for detection of the specific fluorescent reporter. Gating strategies were established using tissues from non-fluorescent wild-type mice to define thresholds for fluorescence-positive populations. Doublets and dead cells were excluded based on forward and side scatter parameters and viability dye staining. Sorted fluorescent-positive cells were collected into tubes containing a collection buffer (e.g., FBS or culture media supplemented with 10–20% FBS) and were either processed immediately for downstream applications or stored at −80 °C or in liquid nitrogen depending on the experimental requirements.

**GeoMx Digital Spatial Profiling**

RNAscope staining was performed using a RNAscope Multiplex Fluorescent Reagent Kit v2 (Advanced Cell Diagnostics), following the manufacturer's protocol. Formalin-fixed paraffin-embedded (FFPE) tissue sections were dried at 60 °C for one hour, deparaffinized, and subjected to a series of pretreatment steps including hydrogen peroxide incubation, heat-mediated antigen retrieval, and protease digestion. Specific probes targeting eGFP (ACD, 538851-C3), Drd1 (ACD, 461901), and Drd2 (ACD, 406501-C2) were hybridized at 40 °C for two hours. Signal amplification was carried out using sequential AMP reagents, and fluorescent signals were visualized with Opal dyes—Opal 520 (Akoya Biosciences, FP1487001KT), Opal 620 (FP1495004KT), and Opal 690 (FP1497001KT). Following hybridization, slides were post-fixed and processed according to NanoString's standard protocol for the GeoMx Digital Spatial Profiler (DSP).

For a spatial transcriptomic analysis, tissue sections were hybridized overnight at 37 °C with probes from the GeoMx™ Mouse Whole Transcriptome Atlas (NanoString). After hybridization, samples were blocked with Buffer W (NanoString) for 30 minutes and stained with a visualization cocktail containing a nuclear dye (SYTO 83, NanoString) for two hours at room temperature in a humidified chamber.

Slides were then imaged on the GeoMx DSP system to generate high-resolution fluorescence maps. Regions of interest (ROIs) were manually selected based on probe signal intensity and tissue architecture, and molecular compartments were defined using fluorescent markers.
Barcoded oligonucleotides from each ROI were photocleaved using UV light and collected via microcapillary aspiration into a 96-well plate. These oligonucleotides were then used for library preparation and sequenced on the Illumina NovaSeq 6000 platform (TheragenBio, Seongnam, Korea) following the manufacturer's instructions.

Raw barcode counts corresponding to RNA targets were analyzed using the GeoMx software suite. Expression values were normalized using the third quartile (Q3) normalization method to account for variability in hybridization efficiency and technical bias across samples.

**Sequencing Data: Processing and Quality Control**

Following spatial transcriptomic profiling with the GeoMx™ Digital Spatial Profiler, raw sequencing data (FASTQ format) were processed using the GeoMx NGS analysis pipeline. Reads were first assessed for quality (Q30), adapter sequences were trimmed, and paired-end reads were merged to produce stitched reads. These were aligned to probe-specific barcode sequences, and PCR duplicates were removed using unique molecular identifiers (UMIs), resulting in deduplicated read counts. The processed data were then imported into the GeoMx DSP Data Analysis Suite for quality control (QC) and downstream interpretation.

QC involved two stages: segment-level QC and biological probe QC. For segment QC, the following thresholds were applied: ≥1,000 raw reads, ≥50% alignment rate, ≥80% stitched and trimmed reads, ≥50% sequencing saturation, geometric mean of negative probe counts ≥1, no-template control counts <2,000, tissue segment area ≥1,000 $\mu m^2$, and ≥50 nuclei per segment. Biological probe QC was performed to exclude underperforming probes. Probes were excluded if their mean count was ≤10% of the total probes for a given gene. Likewise, genes were removed from further analysis if their expression levels fell below the limit of quantitation (LOQ) in more than 20% of segments. The LOQ was calculated as follows: $LOQ = GeoMean (NegProbes) \times [GeoStdev (NegProbes)]^2$ This threshold ensured reliable detection of gene expression above background noise.

Finally, normalization was conducted across samples using the Q3 normalization method. For each segment, the third quartile of expression values was calculated. The geometric mean of Q3 values across all segments was used to derive a normalization factor for each sample. Each expression value was then scaled by this factor, minimizing the influence of outliers and variability in cell density or RNA capture efficiency across segments.

**Multielectrode Array**

Artificial cerebrospinal fluid (ACSF) was prepared with the following components: 92 mM NMDG, 25 mM glucose, 5 mM sodium L-ascorbate, 2.5 mM KCl, 1.25 mM $NaH_2PO_4$, 2 mM thiourea, 10 mM $MgCl_2$, 30 mM $NaHCO_3$, 20 mM HEPES, 3 mM sodium pyruvate, and 0.5 mM $CaCl_2$ (pH adjusted to 7.4). For tissue dissection, a calcium-free variant of ACSF was used by omitting $CaCl_2$.

Mice were deeply anesthetized and transcardially perfused with ice-cold NMDG-based ACSF. Brains were rapidly removed and transferred to chilled, oxygenated NMDG-ACSF. Using a vibratome (VT1000S; Leica Biosystems, Wetzlar, Germany), sagittal brain slices were cut at a thickness of 280 μm. The slices were then incubated in warmed (35 °C), carbogen-bubbled (95% $O_2$/5% $CO_2$) NMDG-ACSF for 30 minutes to allow recovery.

MEA plates (M384-tMEA-6W WHITE; Axion Biosystems) were pre-treated by incubating them in standard ACSF—composed of 92 mM NaCl, 25 mM glucose, 5 mM sodium L-ascorbate, 2.5 mM KCl, 1.25 mM $NaH_2PO_4$, 2 mM thiourea, 2 mM $MgCl_2$, 30 mM $NaHCO_3$, 20 mM HEPES, 3 mM sodium pyruvate, and 2 mM $CaCl_2$—at room temperature for at least one hour.

For recording, individual brain slices were carefully transferred onto the MEA surface and overlaid with recording ACSF containing 124 mM NaCl, 12.5 mM glucose, 2.5 mM KCl, 1.25 mM $NaH_2PO_4$, 2 mM $MgCl_2$, 2 mM $CaCl_2$, 26 mM $NaHCO_3$, and 20 mM HEPES. The MEA plates were then placed into the Maestro Edge system (Axion Biosystems, GA, USA), and spontaneous or optogenetically evoked neural activity was recorded. Electrophysiological data were analyzed using Offline Sorter V4 software (Plexon, TX, USA).

**Optogenetic stimulation**

Optogenetic stimulation was presented using a blue LED plate (170 mm x 130 mm, Scitech Korea Inc., Seoul, Korea) on a MEA plate (Fig. 4b). The optogenetic stimulation was delivered to the sagittal brain slices within the plate through the transparent window of the MEA system where the MEA plate was placed (473 nm, 20 Hz, 50% duty cycle, 10 $mW/Cm^2$ when measured at the bottom of the plate within the MEA system). The optogenetic stimulation consisted of eight cycles of 10s blue light presentation followed by 30-s rest periods. The expression of channelrhodopsin in the MEA NAc was confirmed previously using a confocal microscope (FluoView FV1000, Olympus, Tokyo, Japan).

**Statistical Analysis**

Two-tailed and unpaired or paired t-tests were used in experiments comparing two groups. One-way ANOVA was used for comparing more than two groups and two-way mixed factorial ANOVA was used in experiments requiring between and within group analysis with GraphPad Prism 10. Transcriptome analysis was carried out using gene ontology (GO) analysis in ToppGene (https://toppgene.cchmc.org/), functional annotation clustering analysis in DAVID (https://davidbioinformatics.nih.gov/), and analysis of protein-protein interaction (PPI) in STRING (https://string-db.org/). Graphs of the transcriptome analysis results were produced using Science and Research Online Plots (https://www.bioinformatics.com.cn/srplot). Statistically, p values less than or equal to 0.05 are considered significantly different between conditions. All data are expressed as mean ± SEM (standard error of the mean).

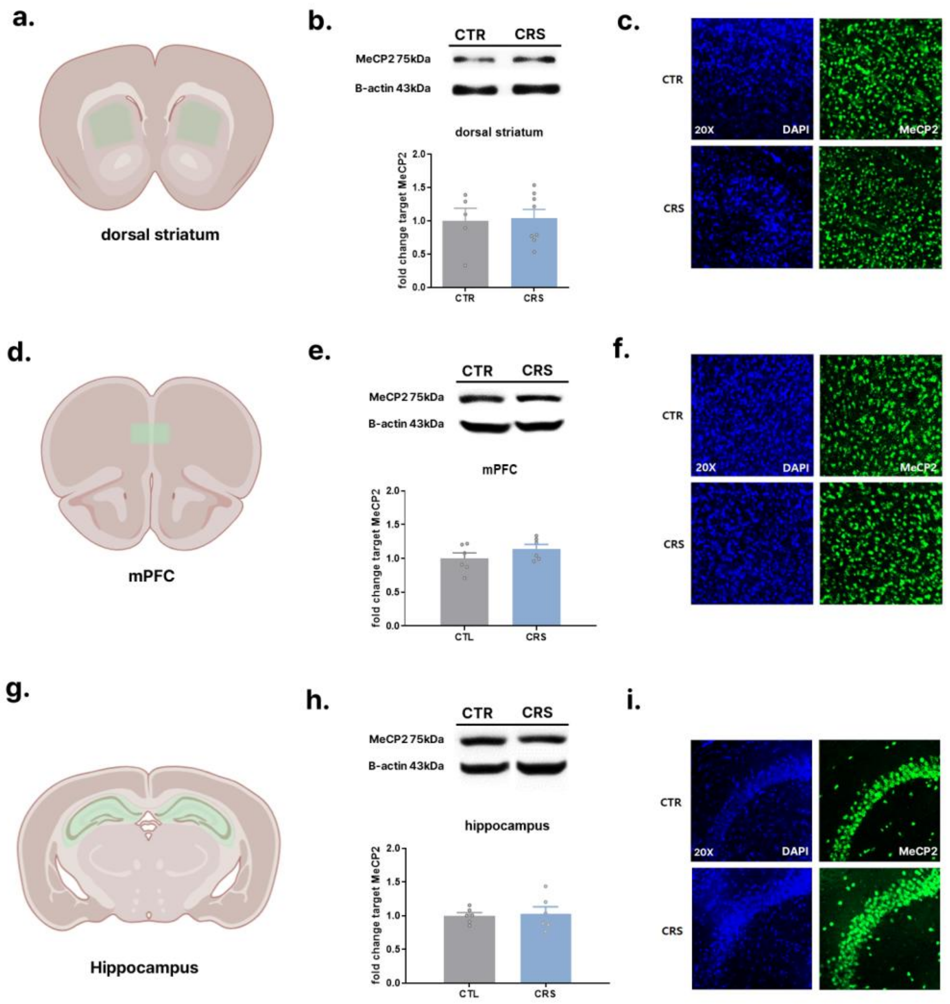


**Supplementary Fig. 1** Confirmation of MeCP2 protein expression in the dorsal striatum, medial prefrontal cortex (mPFC), and hippocampus following chronic restraint stress (CRS). **a–c** MeCP2 expression in the dorsal striatum following CRS. **b** Representative Western blot images (top) and quantification of MeCP2 protein levels normalized to β-actin (bottom); no significant difference was observed between control and CRS groups (two-tailed, t-test, $t = 0.1685$, $p = 0.8692$, $df = 11$, $n = 5, 8$ mice/group). **c** Representative immunofluorescence images of MeCP2 (green) and DAPI (blue) in the dorsal striatum (scale bar: 20× magnification). **d–f** MeCP2 expression in the medial prefrontal cortex (mPFC) following CRS. **e** Representative Western blots (top) and quantification of MeCP2 levels (bottom); no significant difference was observed (two-tailed, t-test, $t = 1.343$, $p = 0.2090$, $df = 10$, $n = 6$ mice/group). **f** Representative immunofluorescence images of MeCP2 and DAPI in the mPFC. **g–i** MeCP2 expression in the hippocampus following CRS. **h** Western blot images and corresponding quantification showing no significant change in MeCP2 expression in the CRS group (two-tailed, t-test, $t = 0.2692$, $p = 0.7933$, $df = 10$, $n = 6$ mice/group). **i** Representative immunofluorescence images of MeCP2 and DAPI in the hippocampus. Data are presented as mean ± SEM (panels b, e, and h).

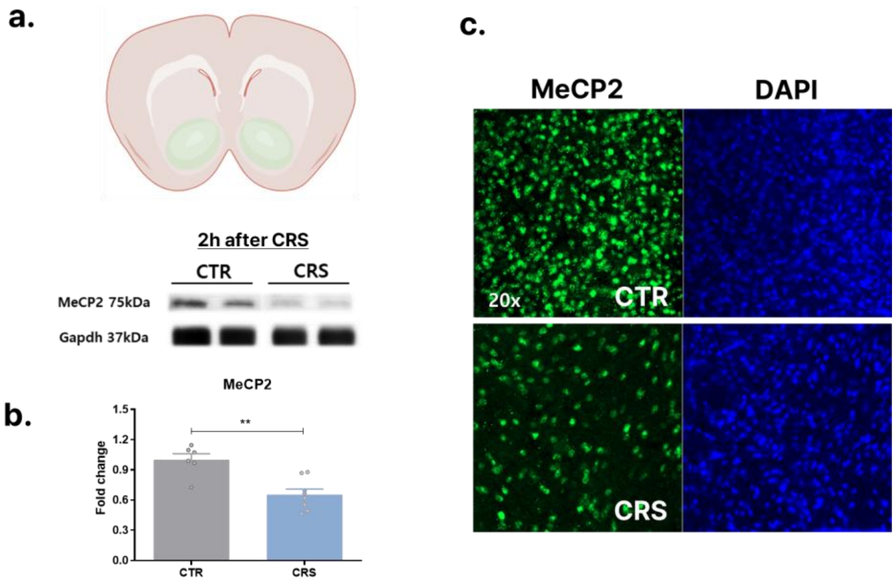


**Supplementary Fig. 2** Reduced expression of MeCP2 protein in NAc by CRS (2 hours after the end of CRS on the last day) **a** Representative immunoblots of NAc **b** Quantification of Western blot bands (two-tailed, t-test, t = 4.166, p = 0.0013, df = 12, n = 6, 8 mice/group), Data are expressed as mean ± SEM, $**p < 0.01$ **c** Representative immunofluorescence image of MeCP2 (20× magnification)

a.

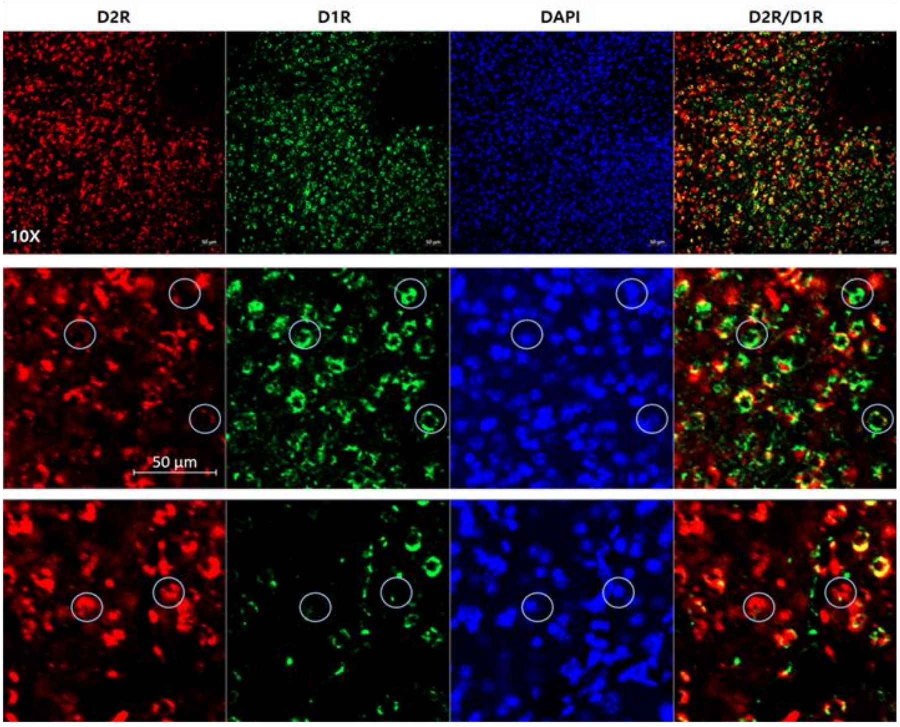
D2R
D1R
DAPI
D2R/D1R
10X
50 µm

b.

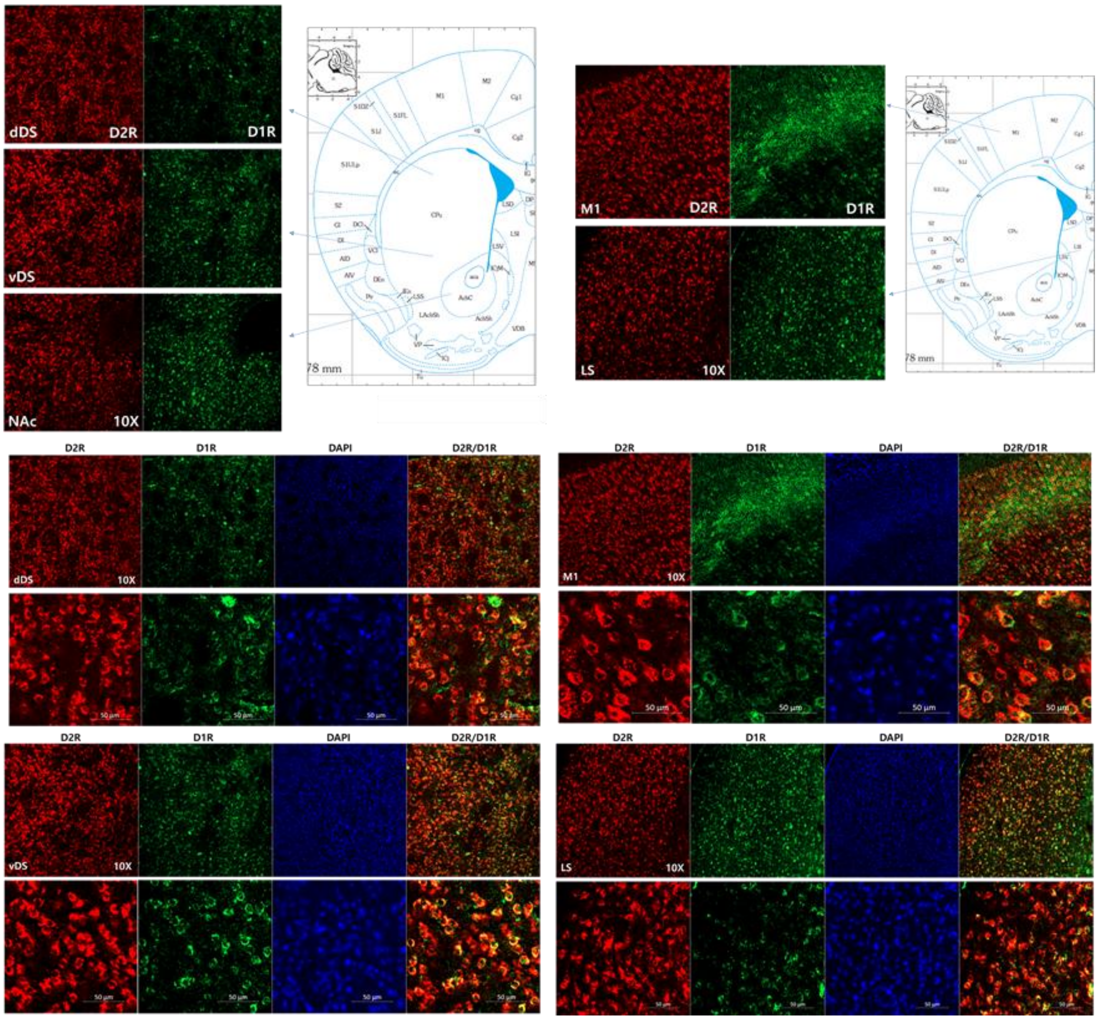
dDS
D2R
D1R
vDS
NAc
10X
M1
LS
DAPI
D2R/D1R
50 µm

**Supplementary Fig. 3 a, b** Representative immunofluorescence images validating the specificity of Drd1 and Drd2 antibodies and confirming the distribution of D1R neurons and D2R neurons. Each specifically labels either the D1 or D2 dopamine receptor. **a** In the NAc, each antibody labels distinct neuronal populations, with a small subset of cells co-expressing both receptors. **b** In other dopamine receptor–expressing regions, D1R and D2R neurons are largely expressed in separate cells, though some overlapping patterns are observed. dDS: upper part of dorsal striatum; vDS: lower part of dorsal striatum; M1: primary motor cortex; LS: lateral septum. Brain atlas reference: "The Mouse Brain in Stereotaxic Coordinates" (Franklin, K. B., 2007)

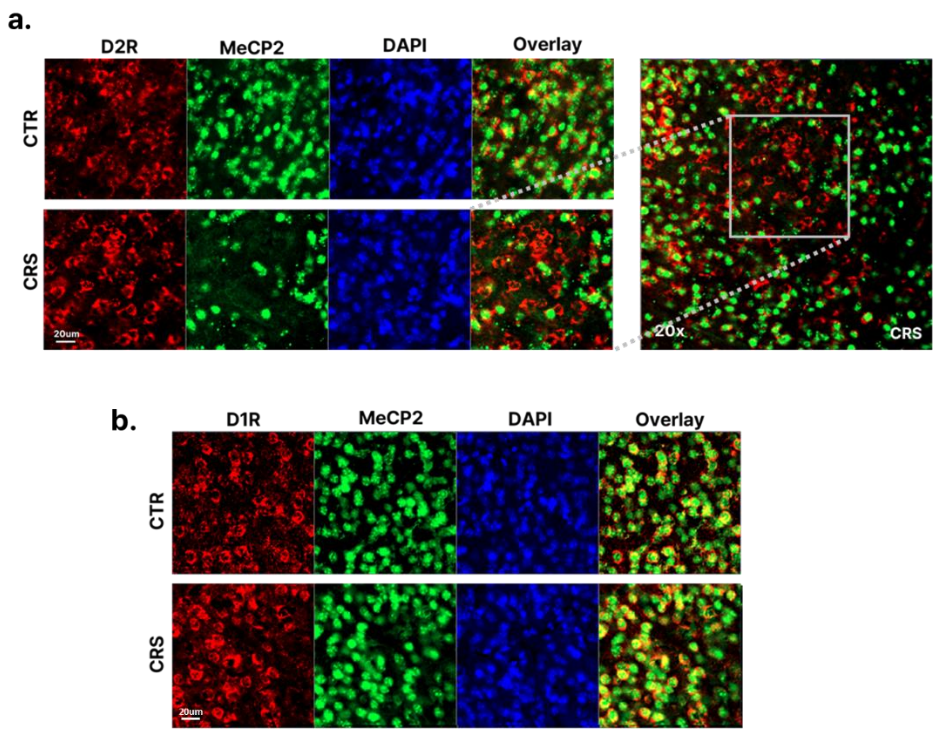


**Supplementary Fig. 4** Representative immunofluorescence images showing cell type–specific expression of MeCP2 in the nucleus accumbens (NAc) 3 days after the final chronic restraint stress (CRS) exposure. **a** A significant decrease in MeCP2 expression was observed in D2R-expressing neurons. **b** No significant change was detected in D1R-expressing neurons. Scale bar: 20 μm. Images were acquired at 20× magnification.

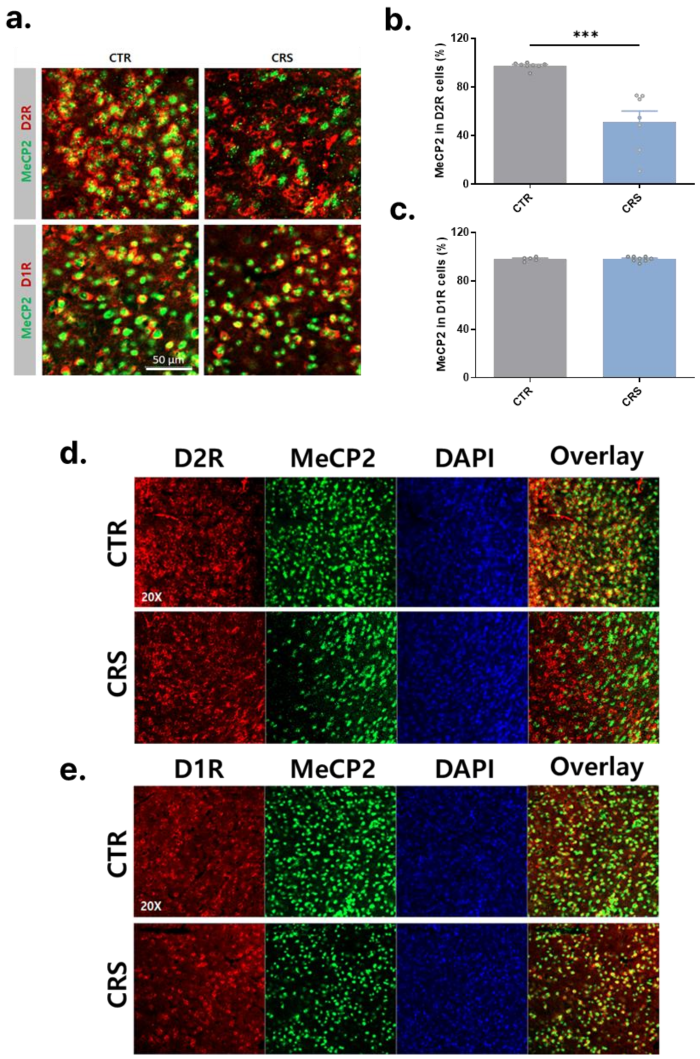


**Supplementary Fig. 5 a-e** Expression of MeCP2 in the NAc by cell type 2 h after the end of the last CRS exposure. **a, d, e** Representative immunofluorescence images showing the expression level of MeCP2 in different cell subtypes (Scale bar 50 μm or 20 X magnification). A significant decrease in MeCP2 was observed in D2 neurons (up), but not in D1R neurons (down). **b, c** Quantification of MeCP2 expression levels by cell type. The proportion of cells expressing MeCP2 in D2R (b) or D1R neurons (c). **b** MeCP2 positive cells in D2R neurons (t-test, two-tailed, $t = 5.447$, $p = 0.0001$, $df = 13$, $n = 8, 7$ mice/group) **c** MeCP2 positive cells in D1R neurons (t-test, two-tailed, $t = 0.045$, $p = 0.\ 0.9651$, $df = 11$, $n = 5, 8$ mice/group). All data are shown as mean ± SEM. ***$p<0.001$

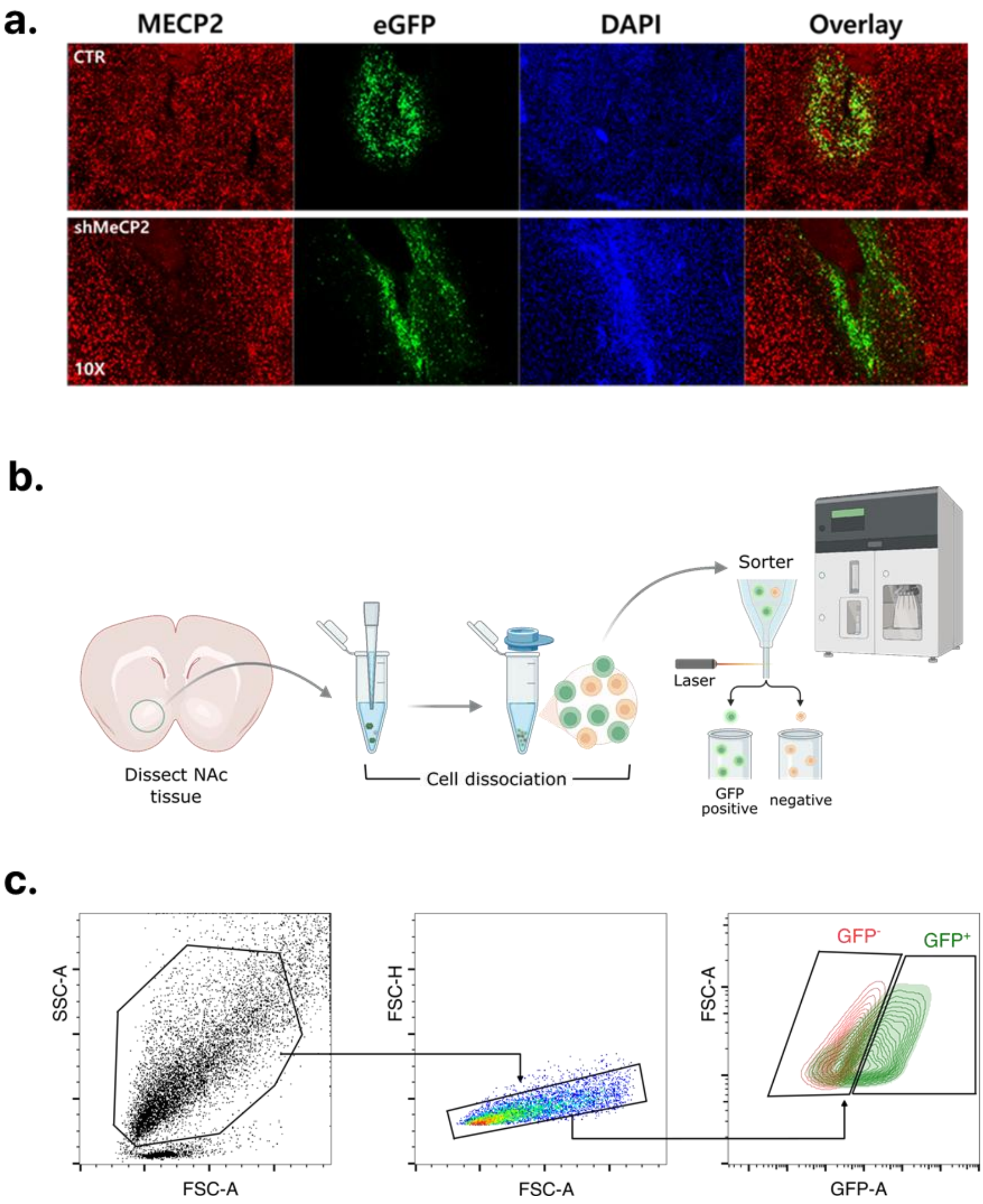


**Supplementary Fig. 6** a Functional validation of AAV- G-CREon-shMeCP2 **a** Representative immunofluorescence images showing reduction of MeCP2 in NAc. Reduction of MeCP2 was confirmed in cells expressing eGFP **b** Illustration of procedure for brain cell dissociation. **c** Gating strategy for fluorescence-based cell sorting in brain tissues.

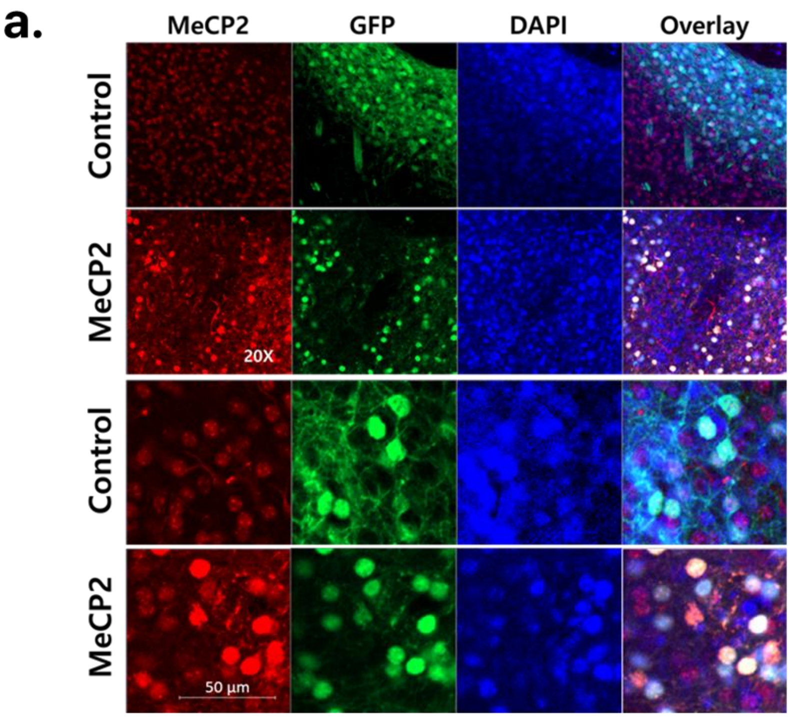


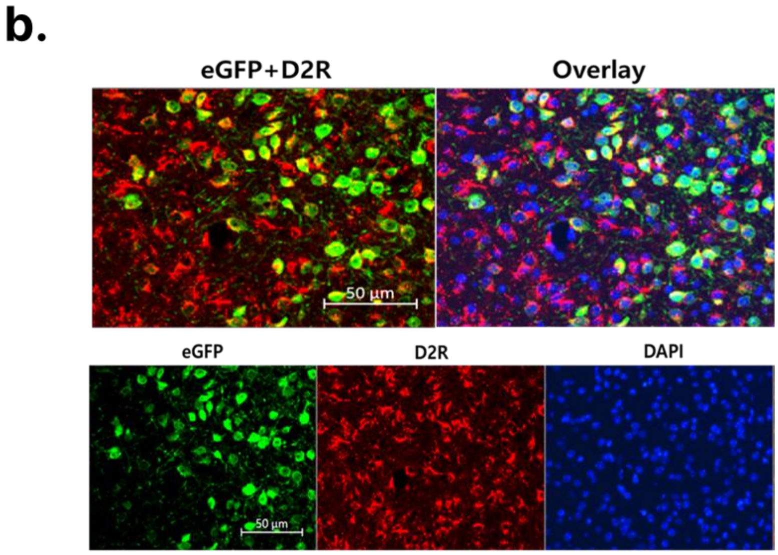


**Supplementary Fig. 7** Representative immunofluorescence images showing increased expression of AAV-MeCP2 in D2R neurons in the NAc by CRE recombinase **a** Increased MeCP2 protein by AAV-MeCP2 in cells coexpressing eGFP and MeCP2. 20x magnification or Scale bar: 50 um **b** Coexpression of eGFP and D2 receptor antibody, showing that AAV is well expressed in target cells. Scale bar: 50 um

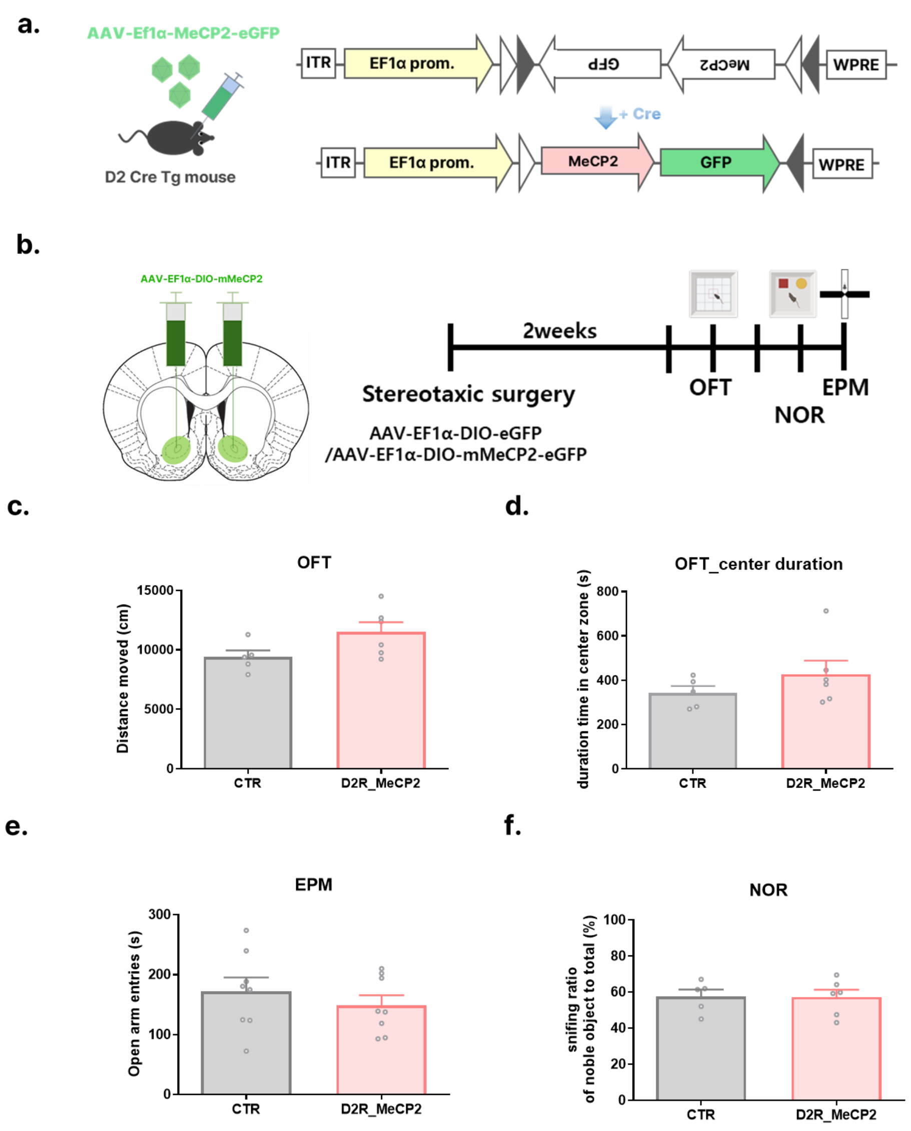


**Supplementary Fig. 8** Effect of genetic MeCP2 increase on D2R neurons in the NAc in normal mice. The augmentation of MeCP2 did not cause abnormalities in motor, anxiety, and cognitive functions. **a** Diagram of AAV-MeCP2 vector expressing mouse MeCP2 mRNA by CRE recombinase b Experimental schedule to confirm the effect of MeCP2 augmentation on D2R neurons in the NAc of normal mice **c** Distance moved in OFT (t-test, two-tailed, $t = 2.013$, $p = 0.0749$, $df = 9$, $n = 5, 6$ mice/group) **d** Time spent in center zone in OFT (t-test, two-tailed, $t = 1.147$, $p = 0.2808$) **e** Time spent in open arms in EPM (t-test, two-tailed, $t = 0.8254$, $p = 0.4230$, $df = 14$, $n = 8, 8$ mice/group) **f** Sniffing time for novel object among total sniffing time in NOR (noble object recognition) test (t-test, two-tailed, $t = 0.047$, $p = 0.9633$, $df = 9$, $n = 5, 6$ mice/group). All data are shown as mean SEM.

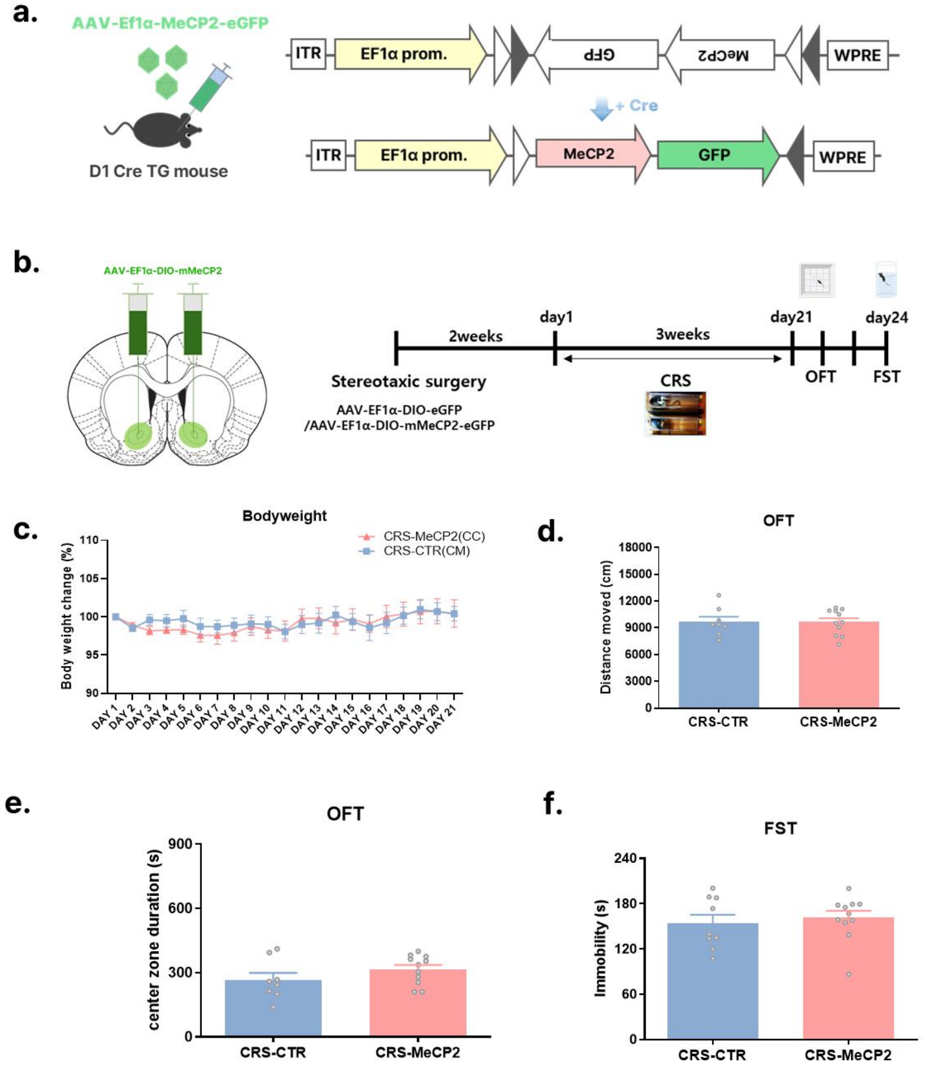


**Supplementary Fig. 9** Effect of MeCP2 regulation of D1R neurons in depression **a** Diagram of vectors that increase the expression of MeCP2 cell type-specifically by Cre recombinase. D1 Cre mice were used to target D1R neurons. **b** Experimental procedures to confirm the effect of MeCP2 increase in D1R neurons in the CRS model. **c** Changes in body weight in the CRS-CTR (control vector) (CC) and CRS-MeCP2 (CM) groups. Two-way Mixed ANOVA, CC vs. CM, $F(1, 17) = 0.041$, $p = 0.8410$ **d** Distance moved in OFT (t-test, two-tailed, $t = 0.026$, $p = 0.9792$, df = 17, n = 8, 11 mice/group) **e** Time spent in the center zone in OFT (t-test, two-tailed, $t = 1.338$, $p = 0.1984$, df = 17) **f** Immobility time in FST (t-test, two-tailed, $t = 0.521$, $p = 0.6084$, df = 18, n = 9, 11 mice/group). All data are shown as mean SEM.

**a.**

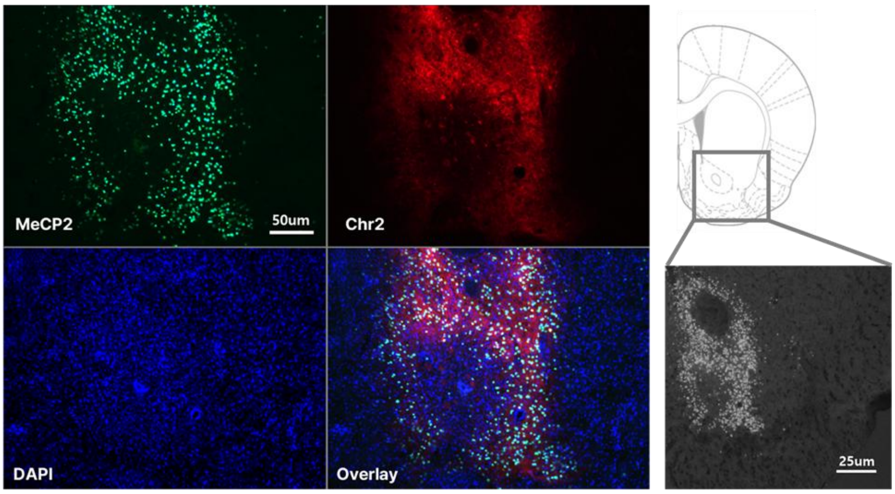


**b.**

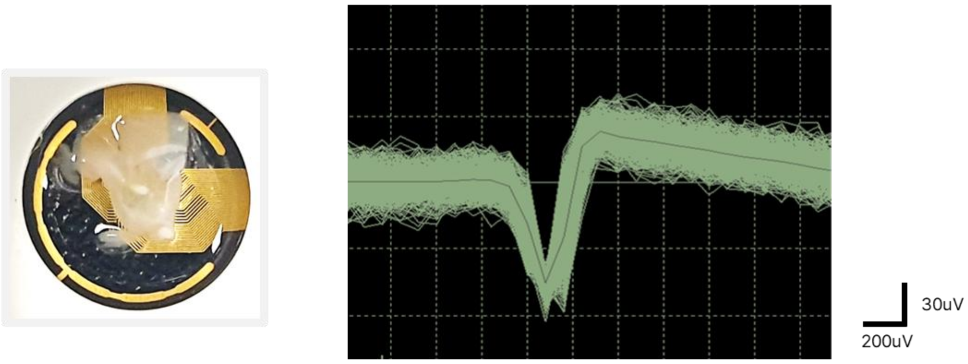


**Supplementary Fig. 10** Recording of neural activity in the NAc using optogenetics and MEA **a** Representative immunofluorescence image showing coexpression of AAV-MeCP2 (green) and AAV-Chr2 (red) in the NAc (left panel). DAPI (blue). Scale bar: 50 µm. Drawing of the mouse NAc indicating the target region and black-and-white image of immunofluorescence showing the area of virus expression (right panel). Scale bar: 25 µm **b** Sagittal section of the brain including the NAc region was placed on a 64-channel MEA recording plate (left panel). Representative spike waveforms generated after optogenetic stimulation in the NAc of the sectioned mouse brain (approximately 250 µm thick).

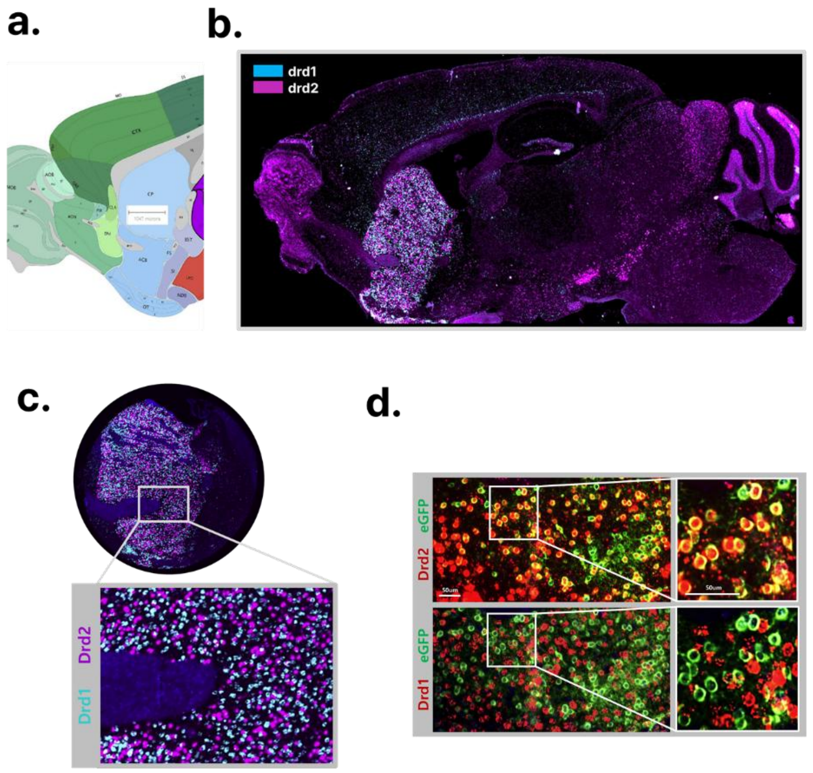


**Supplementary Fig. 11** Cell differentiation using FISH in GeoMX DSP. In the experimental setup for GeoMX transcriptome analysis, FISH was used to visually identify the spatial expression of specific genes within tissues and to select regions of interest for quantitative analysis. **a** Atlas map of a sagittal section of a mouse brain containing the striatum (from Allen brain atlas, https://atlas.brain-map.org/). **b** FISH images to identify cells expressing drd1 and drd2 in the striatum. drd1 (brue), drd2 (purple). The expression levels of drd1 and drd2 mRNA are very high in the striatum. **c** Representative FISH images showing the expression of Drd1 and Drd2 neurons in the NAc. **d** Representative images showing coexpression of Drd2 and eGFP. There is no overlap in the expression of Drd1 and eGFP in the same region, indicating that the AAV was well expressed in the target cells, D2R neurons.

a.

| Target name | Log2 Fold changes | -log10 pvalue |
|---|---|---|
| Adora2a | 21.16 | 9.280 |
| Cre | 16.19 | 9.741 |
| Penk | 15.00 | 12.216 |
| Gpr6 | 11.38 | 11.980 |
| Drd2 | 10.10 | 9.353 |
| Grik3 | 9.58 | 7.147 |
| P2ry1 | 6.84 | 10.177 |
| Gpr52 | 6.60 | 6.988 |
| Unc5d | 4.53 | 5.821 |
| Sp9 | 4.18 | 4.151 |
| Gnas | 3.27 | 5.101 |
| Ecel1 | 3.17 | 1.804 |
| Oprd1 | 3.15 | 5.657 |
| Ptprm | 3.09 | 5.071 |
| Cacna2d2 | 2.99 | 6.491 |
| Sox2 | 2.98 | 5.217 |
| Plxdc1 | 2.78 | 6.481 |
| Htr2c | 2.59 | 6.073 |
| Adk | 2.58 | 4.792 |
| Nrgn | 2.58 | 2.384 |
| Crhr1 | 2.52 | 4.615 |
| Nt5e | 2.49 | 6.071 |
| Tpbgl | 2.40 | 2.626 |
| Fig4 | 2.40 | 4.029 |
| Cbln4 | 2.34 | 3.220 |
| Ndnf | 2.32 | 2.826 |
| Lhx8 | 2.29 | 2.961 |
| Zfp608 | 2.29 | 4.231 |
| Gucy1a1 | 2.25 | 3.786 |
| Sema6c | 2.21 | 4.178 |
| Myo5b | 2.16 | 2.942 |
| Chrna4 | 2.16 | 2.962 |
| B3gat1 | 2.11 | 5.071 |
| Ndst3 | 2.09 | 3.696 |
| Cilp | 2.02 | 1.420 |
| Wnt7a | 2.02 | 2.550 |
| Ebf4 | 2.01 | 2.468 |

c.

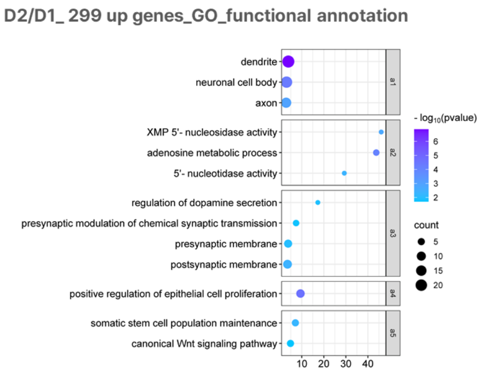


b.

| Target name | Log2 Fold changes | -log10 pvalue |
|---|---|---|
| Ryr1 | -2.00 | 3.352 |
| Prkcg | -2.01 | 3.882 |
| Ltk | -2.04 | 2.986 |
| Nrxn1 | -2.23 | 5.561 |
| Arx | -2.25 | 3.165 |
| Fxyd7 | -2.27 | 3.436 |
| Gnb4 | -2.27 | 5.169 |
| Asic4 | -2.30 | 3.888 |
| Ube2ql1 | -2.41 | 5.167 |
| Cntnap3 | -2.55 | 5.353 |
| Lingo2 | -2.85 | 4.458 |
| Gfra1 | -2.95 | 1.824 |
| Dlk1 | -3.00 | 2.585 |
| Sstr4 | -3.30 | 3.804 |
| Foxp2 | -4.70 | 2.911 |
| Chrm4 | -6.45 | 4.724 |
| Slc35d3 | -7.06 | 5.742 |
| Tac1 | -7.55 | 4.968 |
| Pdyn | -8.82 | 7.503 |
| Drd1 | -9.29 | 7.047 |

d.

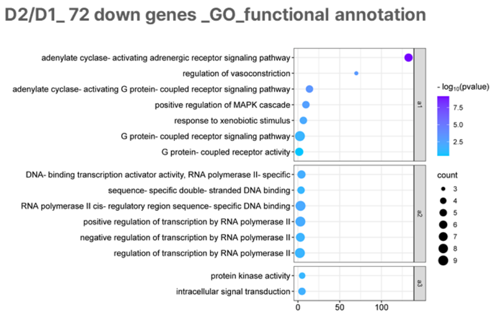


**Supplementary Fig. 12** Comparison of gene expression between D1R and D2R neurons (D2R sample n = 9 mice, D1R sample n = 8 mice). **a** List of genes with higher expression levels in D2R neurons. Drd2 vs Drd1, [FC]>2, -log10 p-values are provided. **b** List of genes with higher expression levels in D1R neurons compared to D2R. [FC] >2, -log10 p-values are provided. -FC values indicate decreased expression relative to D2R neurons. **c, d** Functional cluster analysis of 299 upregulated(c) and 72 downregulated genes(d) in D2R neurons compared to D1R (excluding Cre gene). a1-5: Annotation clusters 1-5, enrichment scores for annotation group >1, p<0.05.

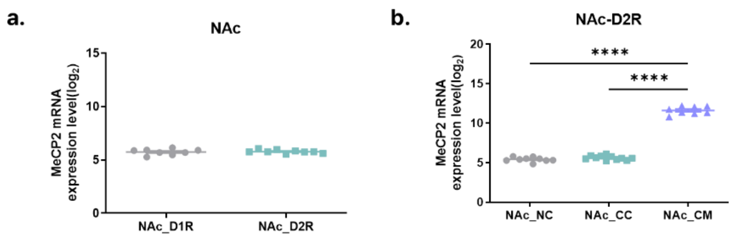


**Supplementary Fig. 13** Comparison of MeCP2 mRNA levels **a** Comparison of mRNA expression rates in D1R and D2R neurons in NAc. t-test, two-tailed, t = 0.3265, p=0.7486, df = 15, n = 8 (D1R samples), 9 (D2R samples) **b** Comparison of MeCP2 mRNA expression levels between groups in NAc D2R samples. NC = naive-CTR (control virus), CC = CRS-CTR, CM = CRS-MeCP2, n = 8, 11, 8 mice/group, One-way ANOVA, F (2, 25) = 820.5, p <0.0001, post-hoc Holm-Šídák's test, NC vs. CC p = 0.2149, NC vs. CM p <0.0001, CC vs. CM p <0.0001, ****p <0.0001, All data are shown as mean SEM.

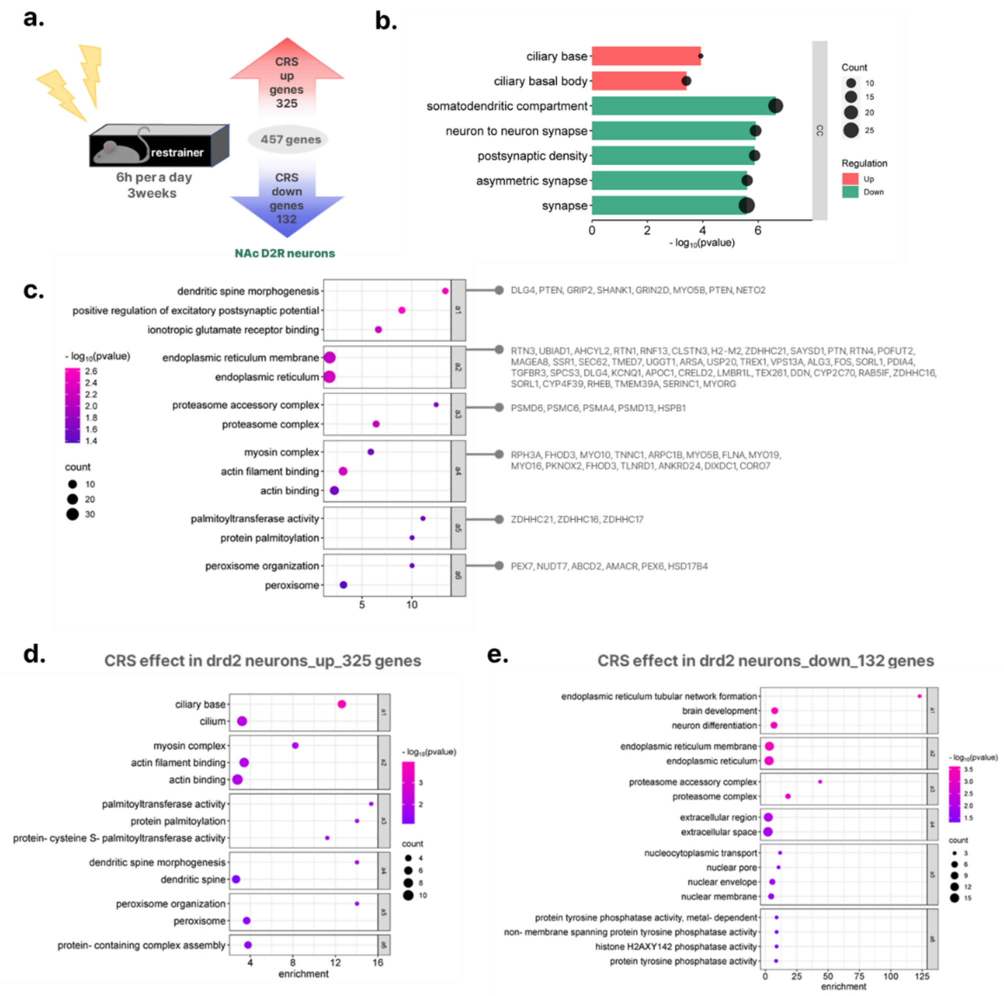


**Supplementary Fig. 14** Effect of CRS on D2R neurons in the NAc. **a** In DEG analysis, 457 genes (up: 325, down: 132) in D2R neurons were changed by CRS ([FC]>1.3, p ≤0.05) n = 8 (NC), 11(CC) mice/group. **b** Gene ontology enrichment analysis for cellular component in NC and CC. Up (red bar) and down (green bar) genes c-e. Functional annotation cluster analysis of 457 genes. We performed DAVID functional annotation cluster analysis from the databases of GO and UniProt (Universal Protein Resource). The three major categories (molecular function, biological process, and cellular component) among the sub-functional categories of GO and Uniprot, respectively, were used as sources for biological function analysis. (c), up-regulated (d), and down-regulated (e) genes whose expression was changed by CRS in D2R neurons of the NAc. The x-axis represents the enrichment score for each function. Enrichment score for annotation groups >1, p <0.05, a1-6 or 1-7: annotation clusters 1-6 or 1-7.

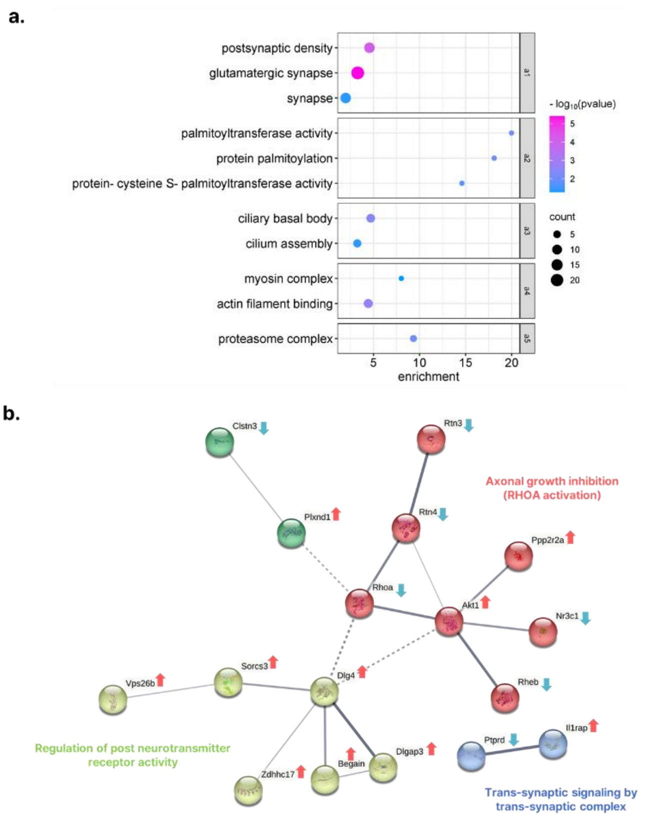


**Supplemental Fig. 15** Genetic MeCP2 increase alleviates gene expression changes induced by CRS (250 genes). CC vs. NC [FC] > 1.3, p ≤0.05; CM vs. NC p >0.10. n = 8 (NC), 11(CC), 8(CM) mice/group **a** DAVID functional annotation clustering graph for total 250 genes, X-axis represents enrichment scores for each cluster. Enrichment score for annotation groups >1, p <0.05, a1-5: annotation clustering 1-4 **c** Functional network analysis (STRING analysis) was performed to show predicted protein-protein interactions for 22 glutamatergic synapse-related genes derived from clustering analysis. Arrows indicate increased (red) or decreased (blue) expression of the corresponding gene by depression. Three functional networks involving glutamatergic synapses were derived.

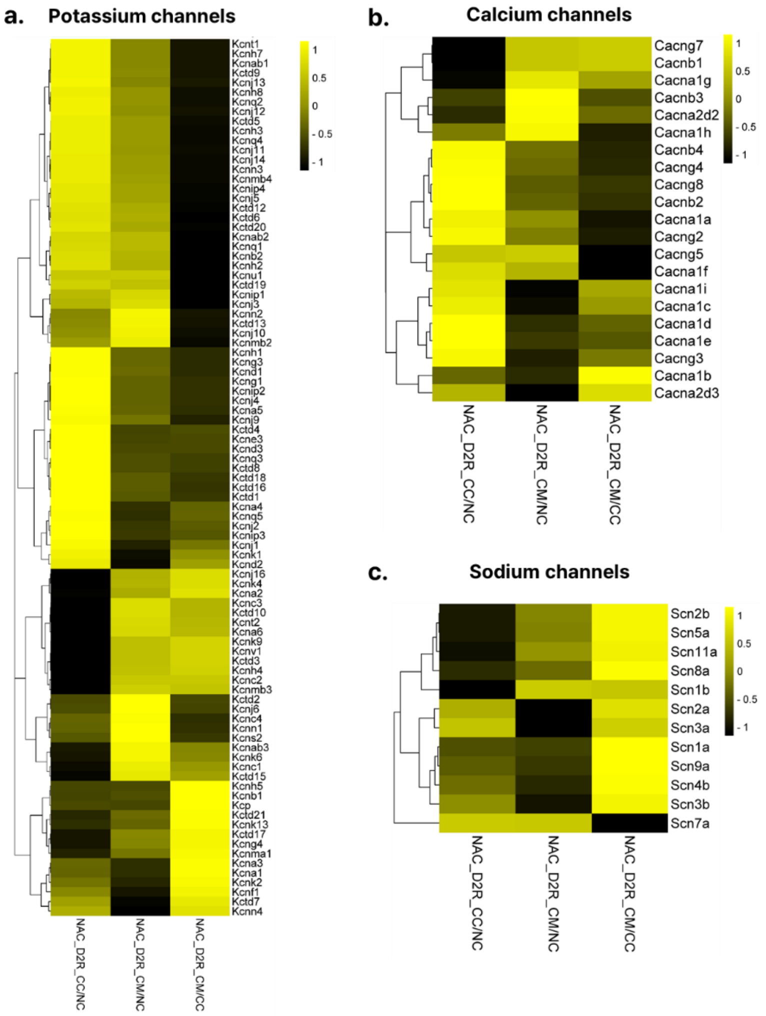


**Supplementary Fig. 16** Heatmap images showing changes in plasticity-related genes by CRS and genetic MeCP2 increase in D2R neurons of the NAc. n = 8 (NC), 11(CC), 8(CM) mice/group **a** potassium channels **b** calcium channels **c** sodium channels, Heatmaps colored from black to yellow according to Z-score scale −1 to 1.

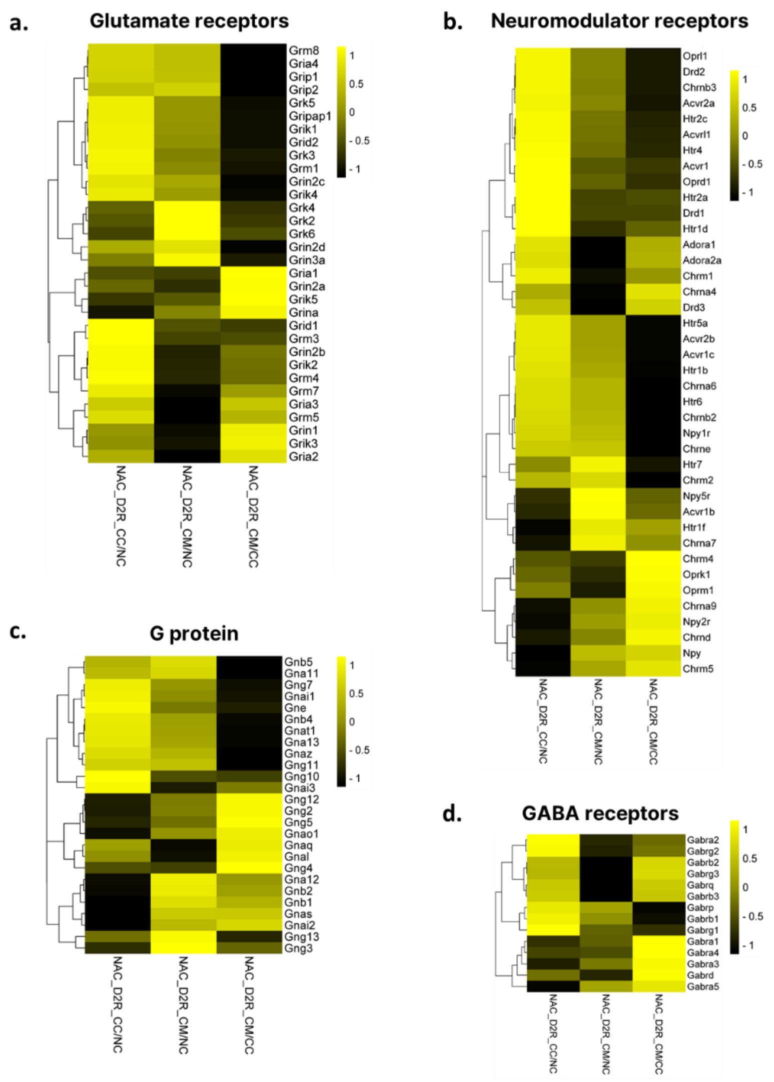


**Supplementary Fig. 17** Heatmap images showing changes in plasticity-related genes by CRS and genetic MeCP2 increase in D2R neurons of the NAc. n = 8 (NC), 11(CC), 8(CM) mice/group **a** Glutamate receptors, **b** neuromodulator receptors, **c** G protein, **d** GABA receptors, Heatmaps colored from black to yellow according to Z-score scale −1 to 1.

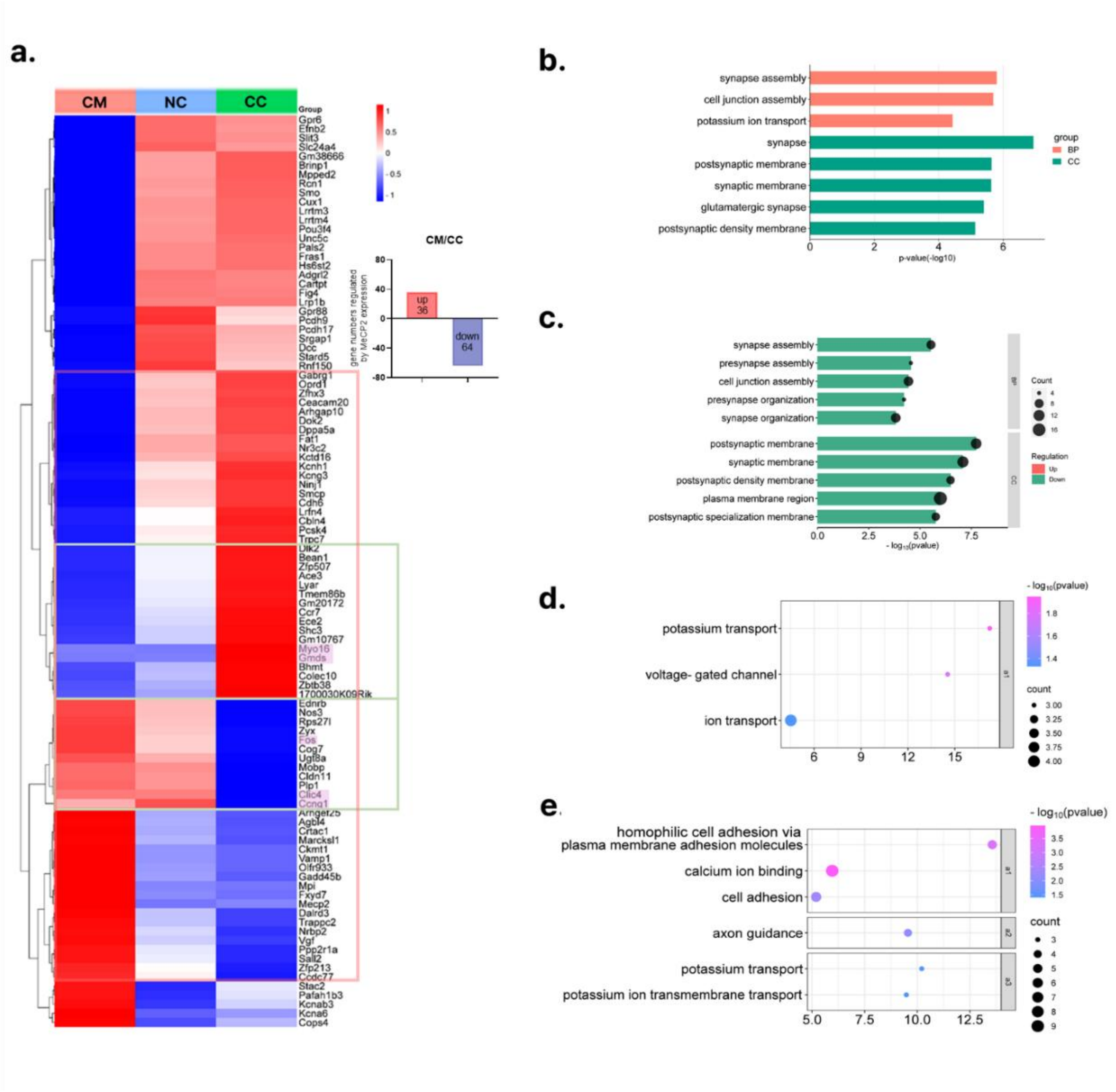


**Supplemental Fig. 18** Effect of genetic MeCP2 increase on D2R neurons in NAc under chronic restraint stress **a** Heatmap image of the relative expression levels of 100 genes (up 36 and down 64 genes) that showed significant differences in CC and CM, CC vs CM [FC] > 1.3, p ≤0.05, pink shades are some of the genes that most significantly recovered the expression levels of genes mutated by CRS by genetic MeCP2 increase (**Fig. 6f, g**). Green boxes indicate areas of clear recovery in the heatmap. **b, c** Gene ontology enrichment analysis for biological process and cellular component in CC and CM. total 100 genes (**b**), up (red bar, no results) and down (green bar) genes (**c**) **d, e** DAVID functional annotation clustering analysis for 100 genes, X-axis represents enrichment scores for each cluster. Enrichment score for annotation groups >1, p <0.05, a1-3: annotation clustering 1-3. Up genes (**d**) and down genes (**e**) were analyzed respectively.

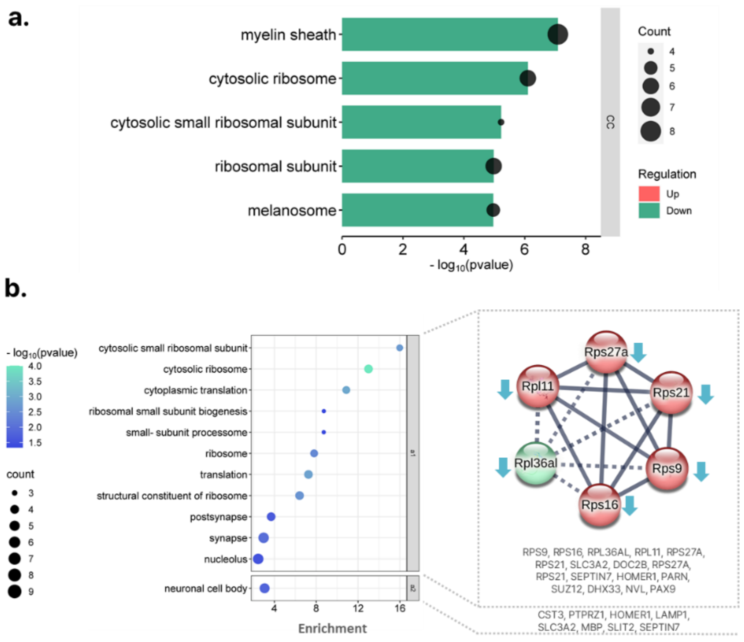


**Supplementary Fig. 19** Effect of CRS in the VP. 122 genes were altered by CRS exposure in VP ([FC]>1.3, p ≤0.05) **a** Enrichment analysis in cellular component of GO for 122 DEGs (FDR <0.05). Top 5 enriched functions are presented. up (red bar, no results) and down genes (green bar). **b** Results of functional annotation cluster analysis after DEG analysis. The X-axis represents the enrichment scores for each function. a1-2: Annotation cluster 1-2 (left panel). Genes corresponding to each main cluster are presented. Image showing the PPI functional network of genes corresponding to annotation cluster 1. The network of the STRING database was used for the analysis

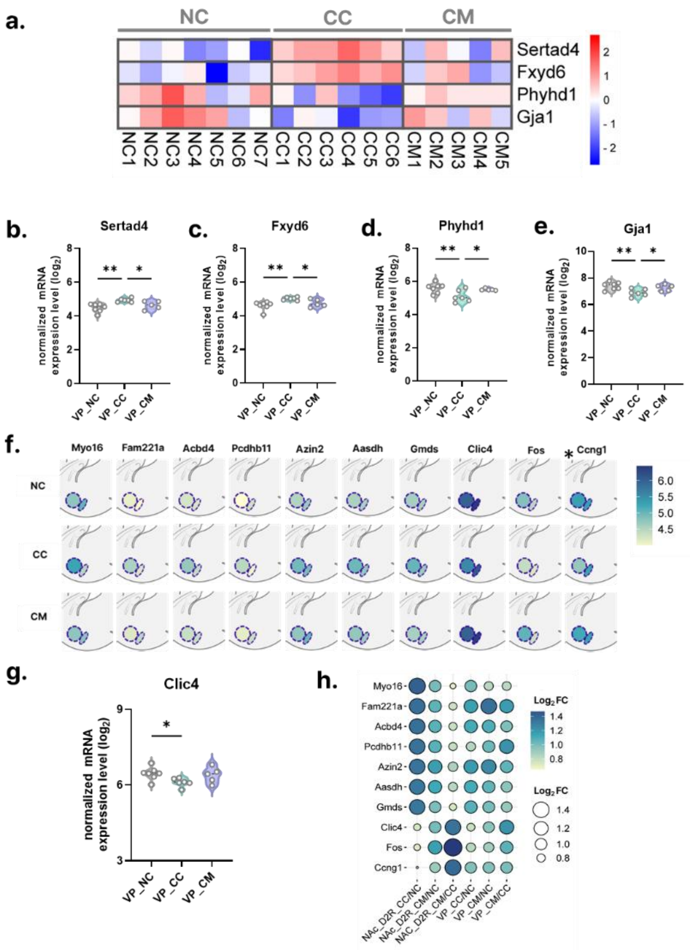

**Supplementary Fig. 20** Circuit-level rescued effect in the VP by genetic MeCP2 increase in NAc. **a-e.** Genes that showed the most significant rescue effect. **a** Heatmap graph showing the expression changes for four genes. Heatmaps colored from blue to red according to Z-score scale **b-e** One-way ANOVA analysis, post-hoc Fisher's test, Sertad4:$F(2, 15) = 7.722$, $p = 0.0049$, NC vs. CC $p = 0.0014$, NC vs. CM $p = 0.2048$, CC vs. CM $p=0.0357$, **c** Fxyd6: $F(2, 15) = 6.459$, $p = 0.0095$, NC vs. CC $p = 0.0028$, NC vs. CM $p = 0.2728$, CC vs. CM $p=0.0462$, **d** Phyhd1: $F(2, 15) = 5.348$, $p = 0.0173$, NC vs. CC $p=0.0066$, NC vs. CM $p=0.5937$, CC vs. CM $p=0.0320$, **e** Gja1: $F(2, 15) = 6.018$, $p = 0.0121$, NC vs. CC $p=0.0041$, NC vs. CM $p=0.4481$, CC vs. CM $p=0.0328$ **f-h** Changes in VP of 10 representative genes showing the remedial effect of MeCP2 in NAc **f** A diagram showing the changes in gene expression between regions and groups. Normalized mRNA expression ($Log_2$) values were used. *Genes showing similar expression changes in VP to NAc. **g** Comparison of expression levels between groups for Clic4 gene. Normalized mRNA expression ($Log_2$) values were used. One-way ANOVA analysis, $F(2, 15) =3.434$, $p = 0.0592$, post-hoc Fisher's test, NC vs. CC $p = 0.0238$, NC vs. CM $p = 0.6679$, CC vs. CM $p= 0.0787$ **h** Bubble plot graph for the comparison of gene expression changes between regions and groups. Log2 FC values were used for the analysis. $*p <0.05$, $**p <0.01$. All data are shown as mean SEM.
